# Topological analysis of neuronal assemblies reveals low-rank structure modulated by cholinergic activity

**DOI:** 10.1101/2025.10.26.684564

**Authors:** Enrique Carlos Arnoldo Hansen, Nicole Sanderson, Sarah Nourin, Virginie Candat, Carina Curto, Germán Sumbre

## Abstract

Neuronal assemblies are fundamental building blocks of brain function (e.g. memory, spatial navigation, eye fixation, etc.). However, the principles underlying the connectivity patterns that support their function have remained elusive. The optic tectum of the zebrafish larva is organized into distinct functional neuronal assemblies. These assemblies display all-or-none preferred activation states and inhibitory competition, mechanisms that improve the decoding of visual information. Here, we combined light-sheet microscopy to capture the dynamics of large neuronal networks (∼2,000 neurons) in the optic tectum; genetic cell-type markers for studying the physiological and functional properties of tectal assemblies; and techniques from topological data analysis to study the dynamic connectivity patterns that enable the emergence and functional role of the neuronal assemblies. We found that during spontaneous activations, tectal assemblies maintain a tight and stable ratio of E-I activity despite the large increase in activity. Topological analysis of the spontaneous activations indicated a low-rank organization of the assemblies and a discrete number of temporal activation patterns. Finally, we observed that the cholinergic system can modulate the topological features of the assemblies to alter their functional role.

## 2 Introduction

Across brain regions, neuronal circuits display different structural architectures, dynamics, and functional roles, typically organized according to neuronal assemblies (groups of highly correlated neurons). Neurons within an assembly work together as a coherent functional unit. Assemblies can be involved in sensory processing, the generation of motor behaviors, or cognitive functions [1, 2, 3, 4]. Coordinated assemblies can lead to precise temporal sequences of neural activity, such as the rhythmic patterns and oscillations of Central Pattern Generators (CPGs) which control repetitive motor behaviors (e.g. walking, breathing, or swimming) [5, 6, 7]. Neuronal assemblies also display attractor-like dynamics; they are based on recurrent connectivity and often display temporal dynamics that settle into stable, self-sustained activity patterns (attractors). This ability to converge to and maintain specific activity states allows these circuits to perform functions such as pattern completion or input discrimination. Attractor-like circuits have been used to model short-term memory [8, 9, 10]; orientation tuning in primary visual cortex [11, 12, 13]; eye fixation [14, 12, 15, 16, 17]; spatial navigation [18, 19, 20, 21]; and have been proposed as a mechanism for faster odor recognition in the olfactory bulb [22]. Recently, experimental evidence for attractor circuits has been observed in the head direction system of drosophila [23], the entorhinal cortex of rodents [24], and somatosensory cortex [25]. Studies of mammalian hypothalamus have also provided evidence of line attractors encoding mating dynamics and an affective internal state [26, 27]. Despite these advances, the principles underlying the connectivity structure mediating the functional role of neuronal assemblies remain elusive.

In zebrafish, the optic tectum plays a key role in detecting the physical properties of visual stimuli (position, size, and direction), processing them, and generating goal-directed motor behaviors. Its ongoing spontaneous activity is organized according to neuronal assemblies. These assemblies mimic the neuronal responses to prey-like visual stimuli, and are organized according to the tectum’s retinotopic map [28, 29, 30]. Their spontaneous activations represent all-or-none preferred network states shaped by mutual inhibition, characteristics reminiscent of winner-take-all dynamics [28]. Using mutant zebrafish larvae with abnormal functional connectivity in the tectum, it was possible to test the effect of the tectal assemblies on visual responses. It was found that the tectal assemblies generate sustained activity and pattern completion, improving stimulus discrimination, visual resolution, and prey-capture behavior [31].

One of the challenges of studying neuronal assemblies is that their structure can be obscured by noise as well as nonlinear features and artifacts of the data. Mathematical methods from algebraic topology allow us to circumvent some of these problems. In recent years, there has been an increased interest in the use of topological data analysis of high-dimensional neural data: (i) topology provides qualitative geometric information, (ii) topological features can be more robust to noise, (iii) topology is a tool for converting local features into global properties, and (iv) topological summaries of data can span a range of values for a parameter, such as a threshold, yielding results that are less sensitive to arbitrary parameter choices [32, 33, 34]. Topological methods can also capture properties of higher-order interactions that cannot be captured by other methods [35], and can be used to compute novel matrix invariants that are unaffected by common nonlinearities [36]. For these reasons, we turned to topological analyses to enhance our understanding of the spatiotemporal functional interactions of neuronal assemblies. Here, we used the zebrafish larva expressing GCaMP6f pan-neuronally in combination with light-sheet microscopy, genetic and immunostaning labeling to determine the neurotransmitter identity of the recorded neurons, and a topological approach to study the structure of neuronal assemblies in the optic tectum. We identified glutamatergic, GABAergic and cholinergic cell types, and found that the tectal assemblies displayed an overrepresentation of excitatory neurons as compared to the overall tectal circuit. During the spontaneous activations of neuronal assemblies, the activity of the GABAergic and glutamatergic populations both increased, but the E-I ratio remained stable. Topological analysis of neuronal correlations within assemblies revealed a low-rank functional connectivity structure reminiscent of attractor circuits. This structure was most pronounced during spontaneous activations. We propose that high-dimensional neural activity in the optic tectum collapses to distinct, low-dimensional subspaces during assembly activations.

The activity of cholinergic neurons was associated with longer spontaneous activations of the assemblies. Since cholinergic neurons are usually associated with attention, we suggest that they increase visual detection by strengthening the functional connectivity of neuronal assemblies involved in spatial visual responses.

## Results

### Neurotransmitter identity of tectal assemblies

To learn about the organizational principles underlying neuronal assemblies in the optic tectum, we first quantified the cell-type composition of the tectal assemblies. In zebrafish larvae, the optic tectum is composed of three cell types according to their neurotransmitter identity: glutamatergic, GABAergic and cholinergic [37, 38, 39, 40]. To identify cell types, we used light-sheet microscopy in combination with double transgenic zebrafish larvae expressing pan-neuronally GCaMP6f and mCherry under a glutamatergic promoter (see Methods). This approach enabled us to monitor the spontaneous activity of the majority of tectal neurons at high acquisition rates (15 Hz), and identify the glutamatergic neurons. GABAergic and the cholinergic neurons were identified using immunostaining following the Ca2+ imaging experiments (Anti-ChAT and Anti-Gad1b antibodies, see Methods). We recorded spontaneous activity of 12 larvae (2169±323 neurons per larva) across the entire optic tectum, in the absence of sensory stimuli. 10 of the 12 larvae were successfully immunostained to be able to distinguish all three cell types: glutamatergic, GABAergic, and cholinergic (Figure 1a).

**Figure 1.**
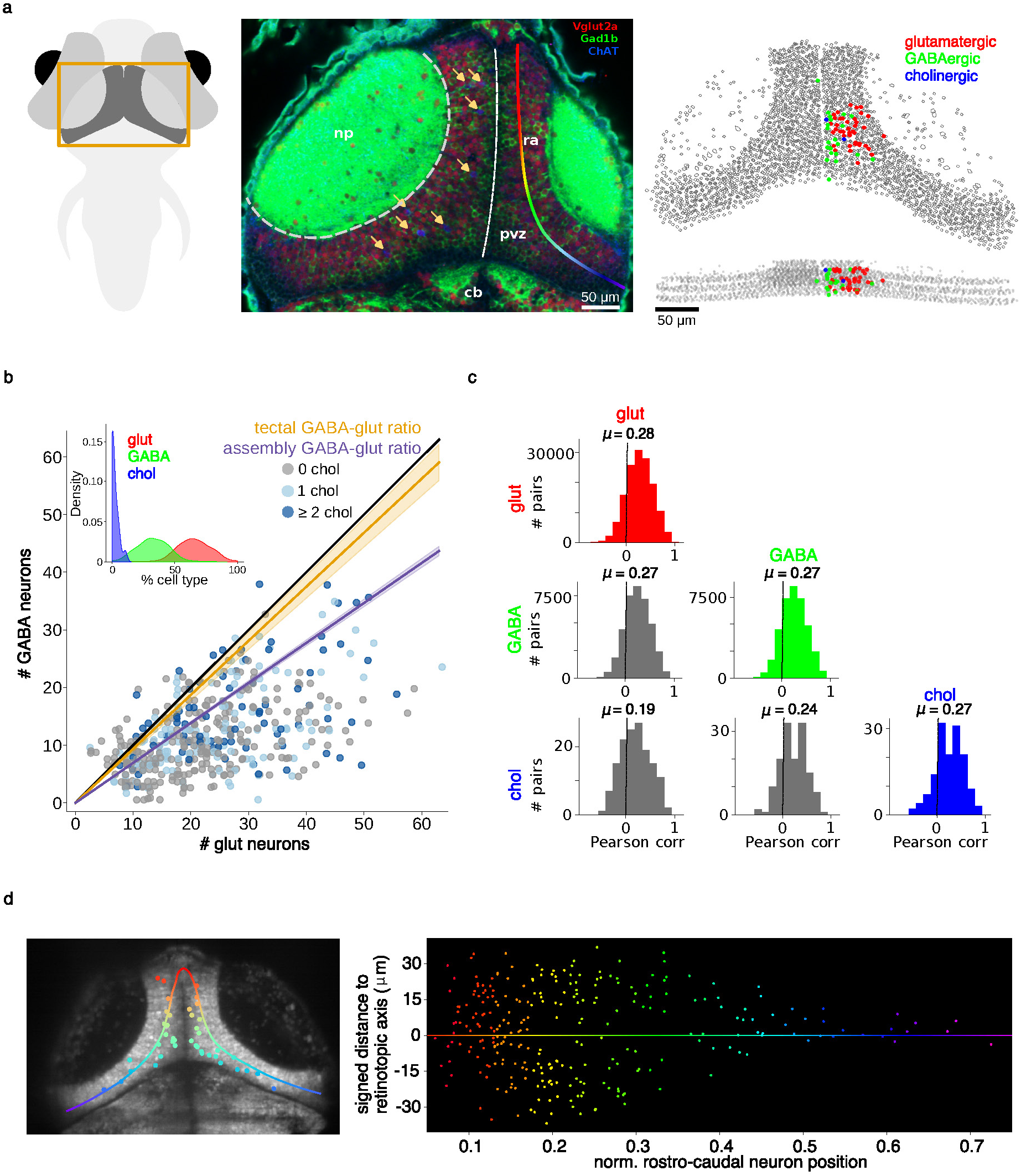
Characterization of neurotransmitter identity of neuronal assemblies in the optic tectum. **(a)** Left: Scheme of the brain of a 6 dfp zebrafish larva. The orange rectangle encloses the optic tectum. Middle: An optical section of the optic tectum following immunochemistry (cb: cerebellum; np: neuropil; pvz: periventricular layer; ra: retinotopic axis). Note the three types of neurons: glutamatergic (red), GABAergic (green), and cholinergic (blue). Yellow arrows point to eight cholinergic neurons. Note their arrangement along the tectum’s retinotopic axis. Right: 3D representation of the tectal neurons. Color: neurotransmitter identity of neurons within one neuronal assembly. Scalebar: 50 µm. **(b)** The numbers of glutamatergic and GABAergic neurons within each of the recorded assemblies. Each dot represents one assembly (*n* = 458 assemblies from 10 larvae). The colors of the dots indicate the number of cholinergic neurons within an assembly. The black diagonal line denotes an equal ratio between glutamatergic and GABAergic populations, while the yellow line shows the expected balance based on the proportion of glutamatergic neurons in the optic tectum (55.47% ± 2.40%, shaded yellow area indicates ± SEM). Note that the assemblies are more glutamatergic, with the purple line indicating the average GABA-glut ratio within assemblies. Cholinergic neurons are scattered across assemblies. Inset: Distributions of the proportions of each population inside the assemblies (glutamatergic: 65.88% ± 13.49%; GABAergic: 32.62% ± 13.13%; cholinergic: 1.93% ± 2.79%). **(c)** Distributions of Pearson’s pairwise correlations between neurons according to their neurotransmitter identity. The mean of the correlations of glutamatergic: 0.28 ± 0.26, GABAergic: 0.27 ± 0.25, cholinergic: 0.27 ± 0.27. The mean correlations between cell types were GABAergic vs. glutamatergic: 0.27 ± 0.27, cholinergic vs. GABAergic: 0.24 ± 0.19, and cholinergic vs. glutamatergic 0.19 ± 0.24. The mean of the entire tectal population correlations: 0.10 ± 0.09 (not shown). **(d)** Left: optical section of the optic tectum showing a projection of all cholinergic neurons (N = 41). Color line: tectal retinotopic axis. Note the cholinergic neurons arranged along the retinotopic axis. Right: the distribution of the cholinergic neurons along the normalized tectal axis from all larvae (N = 278 from 10 larvae).

To detect the neuronal assemblies in the optic tectum, we combined principal component analysis (PCA) and factor analysis (PROMAX) [28, 41, 42, 29] (see Methods). Some assemblies were composed of neurons scattered in both hemispheres of the optic tectum (sparse assemblies) while others were topographically compact, mainly composed of neighboring neurons (compact assemblies) [28]. Altogether, we found 1090 sparse assemblies and 517 compact assemblies in the 10 immunostained larvae. The compact assemblies were identified by examining the distributions of physical distances between neurons in each assembly (see Methods). In this study, we focused only on the compact assemblies (hereon referred as *assemblies*) since their functional role has been previously investigated [28, 43, 29]. These assemblies were composed of 39.39% ± 11.65% of the total recorded neurons, and had an average of 31.54 ± 17.85 neurons.

Quantification of the cell-type composition of the optic tectum showed that the tectal circuit is composed of glutamatergic (55.50% ± 1.65%), GABAergic (43.47% ± 1.78%), and cholinergic (1.31% ± 0.45) neurons (Figure 1b). However, within the assemblies the cell-type distribution was skewed towards excitation: glutamatergic 65.88% ± 13.46%, GABAergic 32.62% ± 13.13%, and cholinergic 1.93% ± 2.79% (see also Supp. Figure 1b-d). Despite the low proportion of cholinergic neurons, they were located along the retinopic axis of the tectum (Figure 1d and Supp. Figure 1a), with none or a few neurons per assembly. Assemblies with cholinergic neurons represented 44.76% of the total number of assemblies.

To learn about the functional connectivity patterns across neurons of different cell types within the neuronal assemblies, we computed Pearson’s pairwise correlations between assembly neurons during spontaneous activations (n=3785 activations of N=438 assemblies). Activations were detected using criteria that capture the fraction of total population activity contributed by the assembly (see Methods). We found that neurons with the same neurotransmitter identity were similarly and highly interconnected (glutamatergic: 0.28 ± 0.26, GABAergic: 0.27 ± 0.25, cholinergic: 0.26 ± 0.27, *P >* 0.05, Mann–Whitney *U* test, Figure 1c). The correlations between the cholinergic neurons and the GABAergic population were not significantly different than those of the GABAergic and glutamatergic ones (0.24 ± 0.26, *P >* 0.05, Krustal-Wallis). However, the correlations between the cholinergic and glutamatergic neurons were significantly lower (0.19 ± 0.29, *P <* 0.05, Krustal-Wallis). The reduced functional correlations between the cholinergic and glutamatergic populations with respect to the other populations may suggest that their interactions are only occasional (e.g. cholinergic modulation of the glutamatergic population).

### Dynamics of different cell types during the activation of tectal assemblies

To further study the functional role of the different cell types during spontaneous activation of the neuronal assemblies, we analyzed the temporal dynamics of the activity of the three cell-type populations. Spontaneous activations occur when a substantial proportion of the neurons in an assembly are active (Figure 2a, top); we detected onsets and offsets of activations using a data-driven algorithm (see Methods). We found that the mean number of active glutamatergic neurons was significantly higher than that of GABAergic neurons, and both were significantly higher than the mean of cholinergic neurons (peak values for glutamatergic: 5.26 ± 2.13, GABAergic: 2.42 ± 1.19, cholinergic: 0.38 ± 0.14; KS tests *P <* 0.0001, Figure 2b).

**Figure 2.**
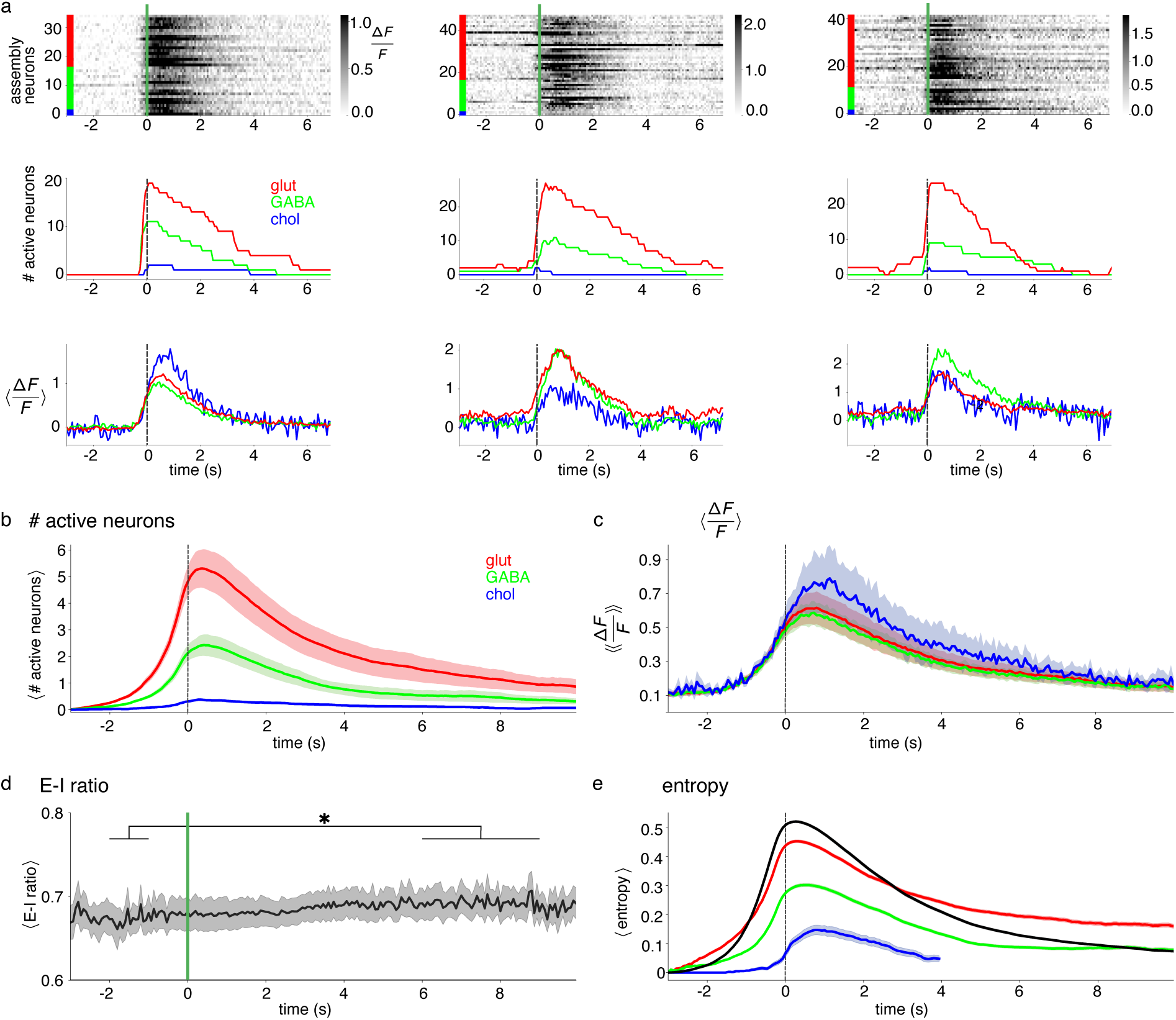
Dynamics of different cell types during spontaneous activation of tectal assemblies. **(a)** Three examples of assembly activations (3 s prior to onset to 7 s after onset), with onsets aligned to time 0. Top: rasters of ΔF/F of the three assemblies (grayscale). Neurons of each assembly are arranged according to their neurotransmitter identity (red: glutamatergic; green: GABAergic; blue: cholinergic). Middle: the number of active neurons in the assembly of each neuronal population. Bottom: average ΔF/F activity of each neuronal population. **(b)** Average number of active neurons (3785 activations of 438 assemblies from 10 larvae). **(c)** Averages of the mean ΔF/F activity across activations for each neuronal cell-type population. Note that the number of active glutamatergic neurons during an activation is higher than that of GABAergic neurons. However, the average activity of the glutamatergic and GABAergic neurons during the time interval under consideration is similar. Although the number of cholinergic neurons is very small, their Ca2+ transients are high. **(d)** The average E-I ratio during the spontaneous activation of the tectal assemblies. Between two and one seconds prior the onset of activations (green line), the value of E-I ratio (0.67 ± 0.06) was lower compared to the value between five to nine seconds after the onset (0.69 ± 0.06). This difference was statistically significant (*P <* 0.001, Kruskal-Wallis followed by Dunn’s test). **(e)** The average entropy of all assembly neurons (black) and the different neuronal cell-type populations (colored curves) during the spontaneous activations of tectal assemblies. Note that the glutamatergic population showed the sharpest increase in entropy prior to the onset of the spontaneous activation of the assemblies, and the highest amplitude. The entropy for the cholinergic neurons was calculated up to 4 s due to the lack of data points. Shaded regions in all panels represent Standard Error Mean.

Despite the larger number of glutamatergic neurons involved in the spontaneous activation of the assemblies, the differences in the mean ΔF/F activity of each cell-type population were not statistically significant (GABAergic: 0.52 ±0.25, glutamatergic: 0.52 ±0.27, and cholinergic: 0.58 ± 0.43; Figure 2c). However, the variability of the cholinergic population was significantly larger than that of the GABAergic and glutamatergic populations (Ansari-Bradley test: *P <* 0.001 for both cases).

We then calculated an excitatory-inhibitory (E-I) ratio, *E/*(*I* + *E*), around the spontaneous activations (see Methods). We found that despite the difference between the number of active excitatory and inhibitory neurons, the E-I ratio remained constant during the entire period of the spontaneous activation of the assemblies (*P >* 0.05, Krustal-Wallis test; Figure 2d). We also observed a slight imbalance between two and one seconds before the onset of activations: the E-I ratio was slightly, but significantly, lower than in the period from five to nine seconds after the onset of activations (before (-2s to-1s): 0.67 ± 0.06, after (5s to 9s): 0.69 ± 0.06, *P <* 0.001, Kruskal-Wallis followed by Dunn’s test).

To further assess the internal structure of assemblies during the spontaneous activations, we computed the normalized entropy for each cell-type population (see Methods). High entropy indicates that the activity patterns are more variable, while low entropy indicates that the repertoire of activity patterns is more limited. We observed that the entropies of the three populations during the assembly activations reached their highest values shortly after the onset of activations, with the glutamatergic neurons having the highest entropy (mean maximal entropy of the glutamatergic neurons: 0.45 ± 0.25; GABAergic neurons: 0.30 ± 0.30; cholinergic neurons: 0.15 ± 0.33; Figure 2e).

Although the spontaneous activations of the neuronal assemblies were dominated by excitation, the E-I ratio remained very stable, consistent with the role of inhibition mainly serving to counterbalance excitation. In contrast, the small number of cholinergic neurons, but with a large and variable ΔF/F, suggests that they may play a modulatory role. The entropy of the spontaneous activations displayed a sharp transition represented by the increase in the number of potential activation patterns (active states). This is mainly evident among the glutamatergic neurons, while the GABAergic population showed a more constrained repertoire of activation patterns. This suggests that although the assemblies are a functional unit, they may be composed of different subgroups of neurons.

To test this hypothesis, we performed community analysis of networks obtained by thresholding assembly correlations to achieve different edge densities (*ρ* = 0.75, 0.5, and 0.25; Supp. Figure 1f). Only in the sparser networks, with *ρ* = 0.25, did we observe significant differences with Erdös-Rényi random graphs (the assembly networks had more communities). This community analysis was not very informative, however, as it was highly sensitive to the choice of thresholds. To avoid this dependency on the choice of threshold, we decided to analyze assembly correlations using topological data analysis (TDA) [34].

### Topological analysis of neuronal assemblies

TDA approaches are sensitive to features such as the higher-order organization of pairwise correlations, going beyond what can be detected using linear algebra tools alone (e.g. PCA, SVD) [44, 35, 45, 24, 34]. In this study we used the clique topology pipeline [36, 46, 34], illustrated here for a small correlation matrix (Figure 3a-c). From a symmetric matrix of pairwise correlations, we obtain a filtration of clique complexes, which are simplicial complexes determined by cliques in an underlying graph. Each clique complex in the filtration has a topological signature given by three integers, (*β*_0_*, β*_1_*, β*_2_), called *Betti numbers*. The resulting topological features across the filtration can be packaged into a collection of *Betti curves*, *β_i_*(*ρ*) (Figure 3c; see Methods). Because the filtration depends only on the relative ordering of the correlation values, the Betti curves are matrix invariants that do not change under nonlinear rescalings of the correlations – provided their relative order is preserved [36]. Therefore, these TDA signatures capture the relative pattern of correlations, independently of their overall magnitudes.

**Figure 3.**
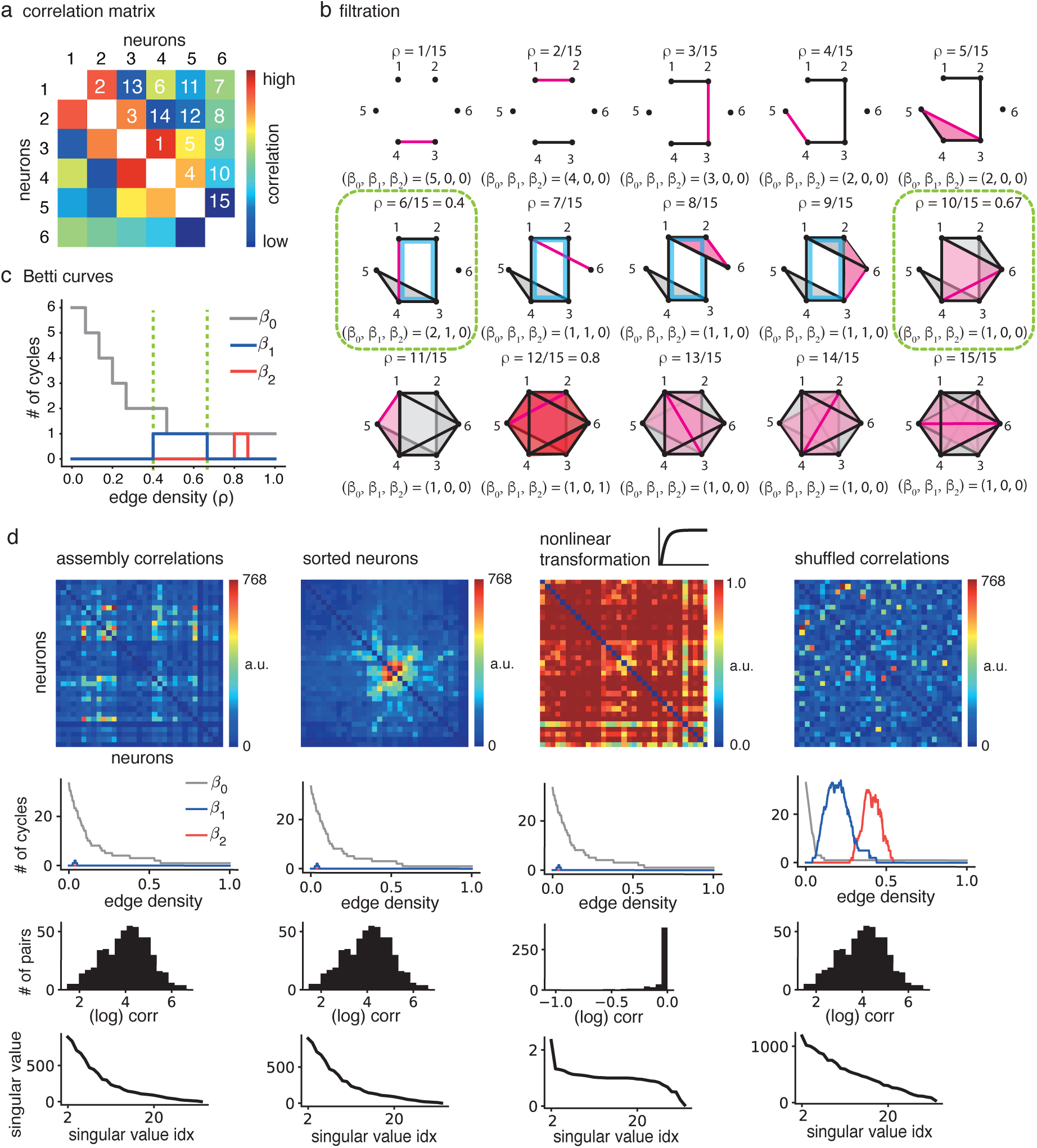
Topological analysis of correlation matrices. We follow the pipeline for computing Betti curves from pairwise correlation matrices given in [36, 34]. **(a)** An example correlation matrix displaying the pairwise correlations of 6 neurons. The white numbers indicate the ranked pairwise correlations from highest to lowest. **(b)** The filtration of simplicial complexes corresponding to the correlations in (a). Each node corresponds to a neuron, and edges are added in a sequence according to their rank, so that pairs of nodes for highly correlated neurons are connected first. One new edge is added at each step of the filtration (shown in pink). Cliques (all-to-all connected subgraphs) are “filled in” to form a simplicial complex from the graph called a clique complex. 2-cliques give rise to edges, 3-cliques are filled in with triangles, 4-cliques with tetrahedra, and so on. These are also shown in pink when they first emerge. A homology 1-cycle can be represented by a cycle in the graph that is not filled in; higher-dimensional cycles are similarly defined (see [34] for details). In this filtration, a homology 1-cycle emerges at step 6 (edge density *ρ* = 6*/*15) and disappears at step 10 (*ρ* = 10*/*15). A homology 2-cycle, corresponding to the 2-dimensional boundary of a void, appears at step 12 and disappears at step 13. Betti numbers (*β*_0_*, β*_1_*, β*_2_) count the number of (non-equivalent) homology cycles in each dimension, with *β*_0_ corresponding to the number of connected components. **(c)** The Betti curves *β_i_*(*ρ*) of the correlation matrix track the Betti numbers at each step in the filtration, parametrized by edge density *ρ*. Notice that *β*_0_ monotonically decreases, as adding edges can never increase the number of connected components. **(d)** (First column) Pairwise correlation for an assembly of neurons, the associated Betti curves, the histogram of correlation values, and the spectrum of singular values. (Second column) Reordering the neurons does not affect the Betti curves nor the distributions. (Third column) Passing the entries of the correlation matrix through a nonlinear, but monotonic, transformation strongly alters the histogram of matrix values and qualitatively changes the singular value spectrum. The Betti curves, however, remain intact. (Fourth column) Shuffling the entries of the correlation matrix leaves the histogram of correlation values intact and produces a milder change in the singular values. However, the Betti curves are completely different and resemble those of a random i.i.d. symmetric matrix.

The Betti curves in Figure 3c illustrate several general phenomena. The *β*_1_ (blue) and *β*_2_ (red) curves always start at zero, may rise for intermediate edge densities, and return to zero by the end of the filtration (*ρ* = 1). In contrast, *β*_0_ (gray) always begins at a positive integer equal to the size of the matrix, and decreases monotonically, as adding edges can only reduce the number of connected components. In this sense, a plateau in *β*_0_ indicates a sub-cluster structure with higher correlations occurring within rather than across connected components (see Supp. Figure 2). Since the complete graph is always connected, *β*_0_ always decreases to 1 by the end of the filtration. In this example, *β*_1_ increases to one with the emergence of the homology 1-cycle (bounding a 2-dimensional hole) at 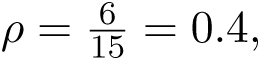 and drops down to zero when the 1-cycle disappears at 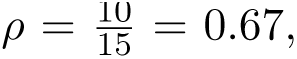 both highlighted with vertical dashed green lines. In contrast, *β*_2_ appears for only one filtration step, at 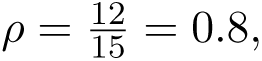 and corresponds to a homology 2-cycle (bounding a 3-dimensional void). When correlations come from time series data, *β*_1_(*ρ*) can reflect sequential dynamics where neurons are activated in a repeating temporal sequence (see Supp. Figure 3), while *β*_2_(*ρ*) can reflect dynamics on a sphere or torus [24]

As mentioned above, Betti curves are distinct from more traditional methods of matrix analysis (e.g., singular values, PCA) because they are invariant to monotone transformations of the correlations while highly sensitive to their higher-order organization [36]. To illustrate this fact, we compared Betti curves to correlation histograms and singular values for four related matrices: a pairwise correlation matrix *C* = [*C_ij_*] of neurons within an assembly, the same matrix with neurons sorted according to hierarchical clustering, a nonlinear monotone transformation of the matrix with entries [*f* (*C_ij_*)] (affecting the values of the correlations but not their relative ordering), and a shuffled matrix containing the same set of pairwise correlations, but scrambling the higher-order structure (Figure 3d, see Methods). For this and all subsequent analyses, we computed pairwise correlations of neurons as simple inner products:

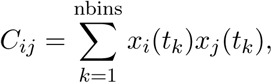

where *x_i_*(*t_k_*) denotes the deconvolved Ca signal of neuron *i* at time *t_k_*, and the sum is computed over different subsets of time bins depending on the analysis (see Methods).

Note that Betti curves, the histogram of correlations, and singular values are all invariant under reordering of neurons in the correlation matrix (Figure 3d, left two columns). However, only the Betti curves are invariant to the nonlinear transformation (third column); the singular values and correlation histograms change significantly. On the other hand, the Betti curves are very sensitive to shuffling the correlations, which changes their organization, while the histograms are not affected by shuffling (fourth column). In summary, Betti curves are uniquely sensitive to the topological organization of the network correlations, rather than to their precise values.

### Betti curves distinguish real versus randomized assemblies

To investigate the structure of the functional correlations underlying tectal assemblies, we computed Betti curves for a large number of assembly correlation matrices. For this analysis, we used all 12 larvae, including the two where immunostaining was unsuccessful, and restricted our analysis to assemblies with at least 20 neurons. We compared the Betti curves of assemblies to those from two families of control matrices: correlations of randomized assemblies and shuffled correlations. Correlation matrices for randomized assemblies were obtained by selecting random subsets of neurons matching the number of neurons in real assemblies; shuffled correlation matrices were obtained as entrywise shuffles of real assembly correlation matrices, as in Figure 3d (see Methods).

First, we looked at *β*_0_ curves, which are indicative of subcluster structure within assemblies. We observed that the average *β*_0_ curves across assemblies (black) and randomized assemblies (yellow) decay more gradually with increasing edge density than that of correlation shuffles (green) (Figure 4a). This suggests that the correlations in the data have a subcluster structure that is not present in the shuffles. Next, we looked at higher-order Betti curves. Unlike *β*_0_, the average *β*_1_ and *β*_2_ curves for the assemblies were very close to zero across all edge densities *ρ* (Figure 4b, left). In contrast, randomized assemblies had higher average *β*_1_ and *β*_2_ curves (Figure 4b, middle), while shuffled correlations exhibited average *β*_1_ and *β*_2_ curves that were orders of magnitude higher than those from the data (Figure 4b, right). These trends in average Betti curves across real versus randomized assemblies are consistent across larvae (see Supp. Figure 4). Thus, Betti curves allow us to detect the functional connectivity structure of assemblies.

**Figure 4.**
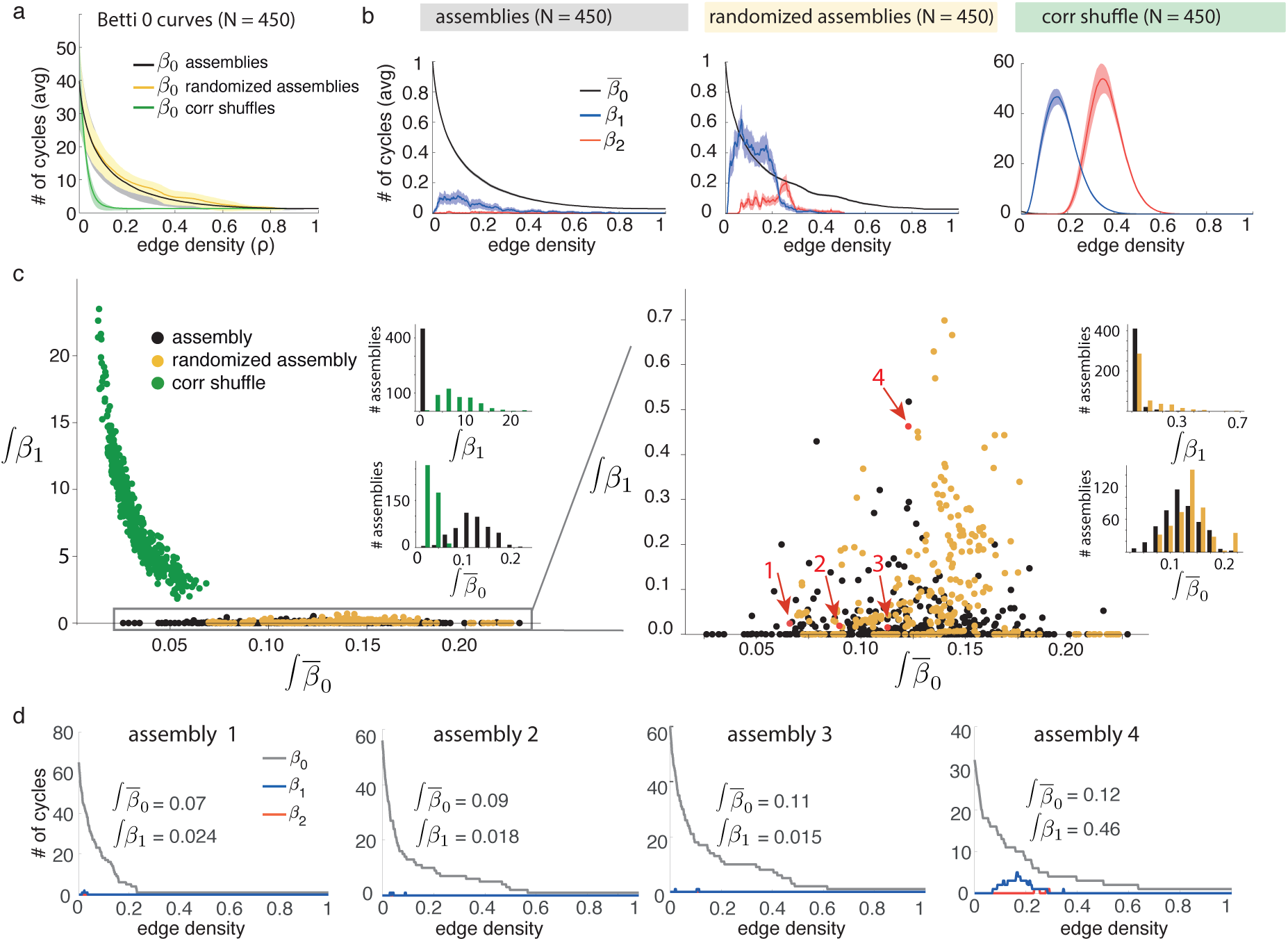
Betti curves capture the special correlation structure of tectal assemblies. **(a)** Average *β*_0_ curves for pairwise correlations between the neurons within the tectal assemblies (black), randomized assemblies (yellow) and entrywise shuffles of assembly correlations (green). Note that shuffling the correlations results in a fast decaying *β*_0_ (N = 450 for 12 larvae). **(b)** Left: Average *β*_1_ curves (blue) and *β*_2_ curves (red) for assembly correlations indicate few homology 1-and 2-cycles; average 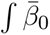 curves are normalized (black). Middle: Betti curves for random assem-bly correlations display more homology 1-and 2-cycles. Right: entrywise shuffles of assembly correlations have large average *β*_1_ and *β*_2_ curves. Notice the increase in the scale of the y-axis. **(c)** Left: The integrated Betti values 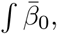 and ∫*β*_1_ from (a) and (b). Assemblies (black) and randomized assemblies (yellow) have much lower ∫ *β*_1_, but higher 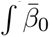 than correlation shuffles (green). Inset: histograms of the distributions of ∫ *β*_1_ (top) and 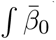 (bottom). Right: zoom of randomized assemblies (yellow). **(d)** Betti curves for individual assemblies annotated in (c, red arrows). Note that 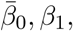 increases from assembly 1 to 4, capturing the slower decay of *β*_0_.

To quantify the above observations, we calculated the *integrated Betti values*, ∫ *β_i_*, corresponding to the areas under the Betti curves [36] (see Methods). The integrated Betti values capture features of Betti curves. For example, if *β*_0_ decays to 1 faster, ∫ *β*_0_ will be smaller. However, the integrated Betti values may have some effects due to the sizes of the matrices. Thus, it is important that the assembly, randomized assembly, and shuffled correlation matrix families have matching distributions of sizes (see Supp. Figure 5-6). To account for specific effects of assembly size on *β*_0_, we used a normalized integrated Betti 0 value, 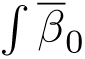 (see Methods). This normalization allows us to more fairly compare the decay rates of *β*_0_ in order to assess subcluster structure. Notice that in Figure 4d, *β*_0_ decays as a function of edge density more slowly as we move from assembly 1 to assembly 4 (left to right); this is reflected in the increase in 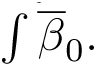.

Consistent with our observations from Figure 4a, the distribution of 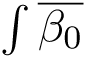 across assembly correlations was significantly different from those of shuffled controls (assemblies: 0.121 ± 0.035, randomized assemblies: 0.138 ± 0.035, shuffled corr. matrices: 0.034 ± 0.011, *P* ≪ 0.001, two-sample two-way KS-test, Figure 4c, left). Moreover, the distribution of ∫ *β*_1_ for assemblies and randomized assemblies were significantly different (assemblies: 0.023 ± 0.09, randomized assemblies: 0.092 ± 0.123, *P* ≪ 0.001, two-sample two-way KS-test, Figure 4c, right). We did not compare ∫ *β*_2_ as it was zero for the majority of assemblies. We conclude that the Betti curves *β*_0_(*ρ*) and *β*_1_(*ρ*) can distinguish the structure of assembly correlations from the controls.

### Spontaneous activations of assemblies exhibit low-rank structure

To study the topological structure of the correlations of tectal assemblies, we analyzed a total of 4595 spontaneous activations across 435 assemblies with at least 20 neurons and at least 2 activations (see Methods, Figure 5a). On average, these assemblies had 10.6 ± 8.5 spontaneous activations, and were active for a total of 26.5 ± 22.6 s across the entire hour-long recordings. The average duration of these spontaneous activations was 2.8 ± 1.8 s.

**Figure 5.**
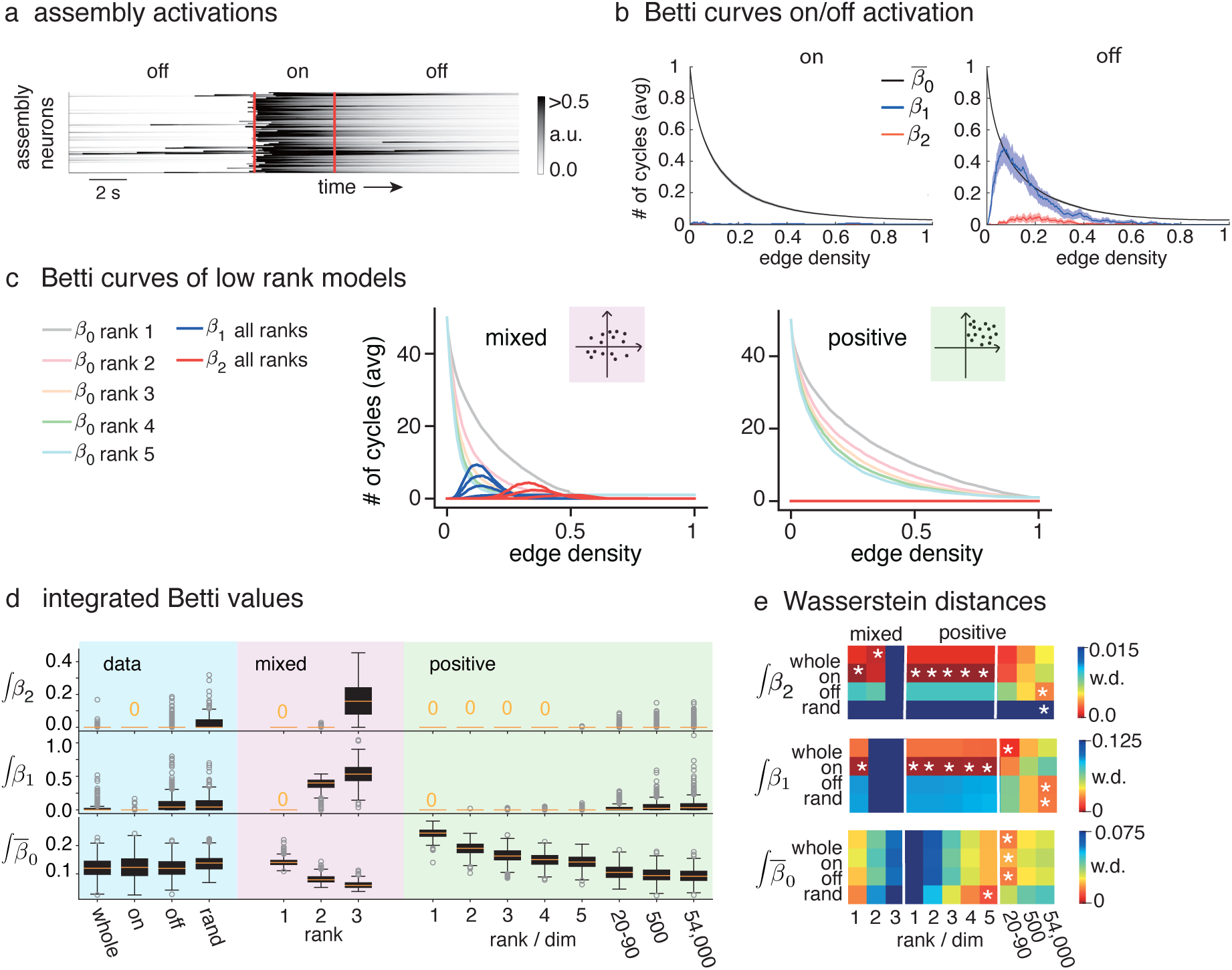
Tectal assemblies display topological features associated with low-rank models. **(a)** An example rasterplot for the neurons within an assembly around a spontaneous activation. The ‘on’ period lies between the activation onset and offset (red lines), the rest is the ‘off’ period. **(b)** Average 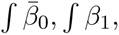 and *β*_2_ curves for assembly correlations during ‘on’ and ‘off’ periods (N = 435 for 12 larvae). Note that *β*_1_ and *β*_2_ are negligible during the on periods. **(c)** Average Betti curves for 100 symmetric matrices of size 50 from two families of low rank models: ‘mixed’ and ‘positive’ low rank matrices (see Methods). *β*_0_ curves for ranks 1-5 (pastel colors) decay more quickly for higher ranks. Note that *β*_1_ (blue) and *β*_2_ (red) curves are unimodal for the ‘mixed’ models, but vanish for the ‘positive’ models. **(d)** Boxplots of the distributions of 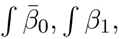 and *β*_2_ for the experimental data, low-rank ‘mixed’ models, and ‘positive’ models of different ranks/dimensions (N=450 matrices of size 50 for each model). **(e)** Wasserstein distances between the 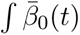 and ∫ *β*_2_ distributions for the experimental data (rows) and the models (columns). White asterisks indicate the closest models for each set of data-derived correlation matrices (‘whole’, ‘on’, ‘off’, and ‘rand’). Note that the assembly correlations during ‘on’ periods best match ∫ *β*_1_ and ∫ *β*_2_ values obtained for the ‘mixed’ rank 1 and ‘positive’ low-rank models. In contrast, the best matches for the ‘off’ and randomized assembly correlations are the models with the highest ranks/dimensions.

To study the structure of the assemblies during periods of spontaneous activation (‘on’) and non-activation (‘off’), we compared the Betti curves during the ‘on’ and ‘off’ periods (see Figure 5b, see Methods). We observed only a small significant difference between the distributions of 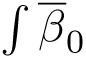 for the ‘on’ periods versus ‘off’ periods, (‘on’: 0.12423± 0.04137, ‘off’: 0.12135 ± 0.03238, P ∼ 0.01, two-sample two-way KS test). However, the ∫ *β*_1_ for the ‘on’ periods were significantly smaller than those from the ‘off’ periods (‘on’: 0.00145 ± 0.01094, ‘off’: 0.09246 ± 0.13060, *P* ≪ 0.001, two sample KS-test).

Next, we investigated whether the Betti curves were indicative of low-rank structure, and thus low-dimensional dynamics. Because of the nonlinearities in data acquisition and preprocessing, this structure cannot be seen via traditional linear algebra methods such as singular values (Supp. Figure 7). Recently, it was proven that for positive symmetric matrices of rank one, all Betti curves vanish except for *β*_0_(*ρ*) [47]. To understand what other types of correlation structures Betti curves can detect, we computed Betti curves for different types of matrices. Specifically, we considered two classes of random matrix models: ‘mixed’ and ‘positive’ matrices for low and high rank controls (see Methods). The ‘mixed’ model is the most standard (default) model for generating random low rank matrices; however, the ‘positive’ model, with matrices generated from random vectors whose entries are all positive, is in principle a better control for our calcium imaging data since our time series values were always nonnegative. Additional analyses further confirmed that the ‘positive’ model is the better control family (see Supp. Figure 8).

The ‘mixed’ models had non-trivial *β*_1_ and *β*_2_ curves at the higher ranks while *β*_0_ displayed some variability (Figure 5c, left). In contrast, ‘positive’ matrices had *β*_1_ and *β*_2_ curves that vanished in each rank (Figure 5c, right). To further assess the structure of the assembly correlations identified by the Betti curves, we calculated the distributions of integrated Betti values of the assemblies for the ‘whole’ recording, during the ‘on’ periods, during the ‘off’ periods, and for randomized assemblies (‘rand’). We compared these distributions against those of synthetic ‘mixed’ and ‘positive’ matrix models of varying dimensions or ranks (Figure 5d). In addition to low ranks, we considered three high-dimensional controls (see Methods): an ensemble of ranks 20-90 (the sizes of the neural assemblies), a model of underlying dimension 500 (the approximate number of time bins for all ‘on’ periods), and another of dimension 54,000 (the total number of time bins in the recording). These last two models produce correlations coming from random time series of equal lengths to those used to compute the assembly correlations. To compare the experimental observations to the different models, we then computed the Wasserstein distances between the distributions for the data correlations and models (Figure 5e). We found that 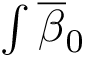 was not informative about the structure of the correlations in different contexts (‘whole’, ‘on’, ‘off’, ‘rand’). However, using ∫ *β*_1_ and ∫ *β*_2_ it was possible to separate the different contexts.

The ‘whole’ and ‘on’ activations showed ∫β_1_ and ∫β_1_ similar to those for both the ‘mixed’ and ‘positive’ low-rank models. In contrast, the distributions of *β*_1_ and *β*_2_ for the ‘off’ and ‘rand’ contexts were only similar to high rank positive models.

These results suggest that during the activation periods (‘on’), the assembly functional connectivity reflects an especially low-rank structure, while between activations (‘off’) the structure is similar to that of randomized assemblies. Despite the fact that the ‘on’ periods are a small fraction of the total length of the recording, the structure of correlations computed from the entire recording (‘whole’) more closely resembled that of the ‘on’ rather than the ‘off’ periods. This indicates that there was little contribution to the structure of assembly correlations outside the activation periods. Note that traditional singular value analyses did not reflect low rank structure during spontaneous activation (Supp. Figure 7).

### Dynamics of the topological structure of the assemblies

To investigate changes in the structure of correlations around the onset of activations, we combined the clique topology pipeline with a sliding window analysis. Specifically, we computed the integrated Betti values 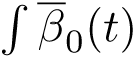 and *β*_1_(*t*) for correlations using a 0.5 s causal sliding window that runs from 2s before to 2s after the onset of each spontaneous activation (Figure 6a, see Methods).

**Figure 6.**
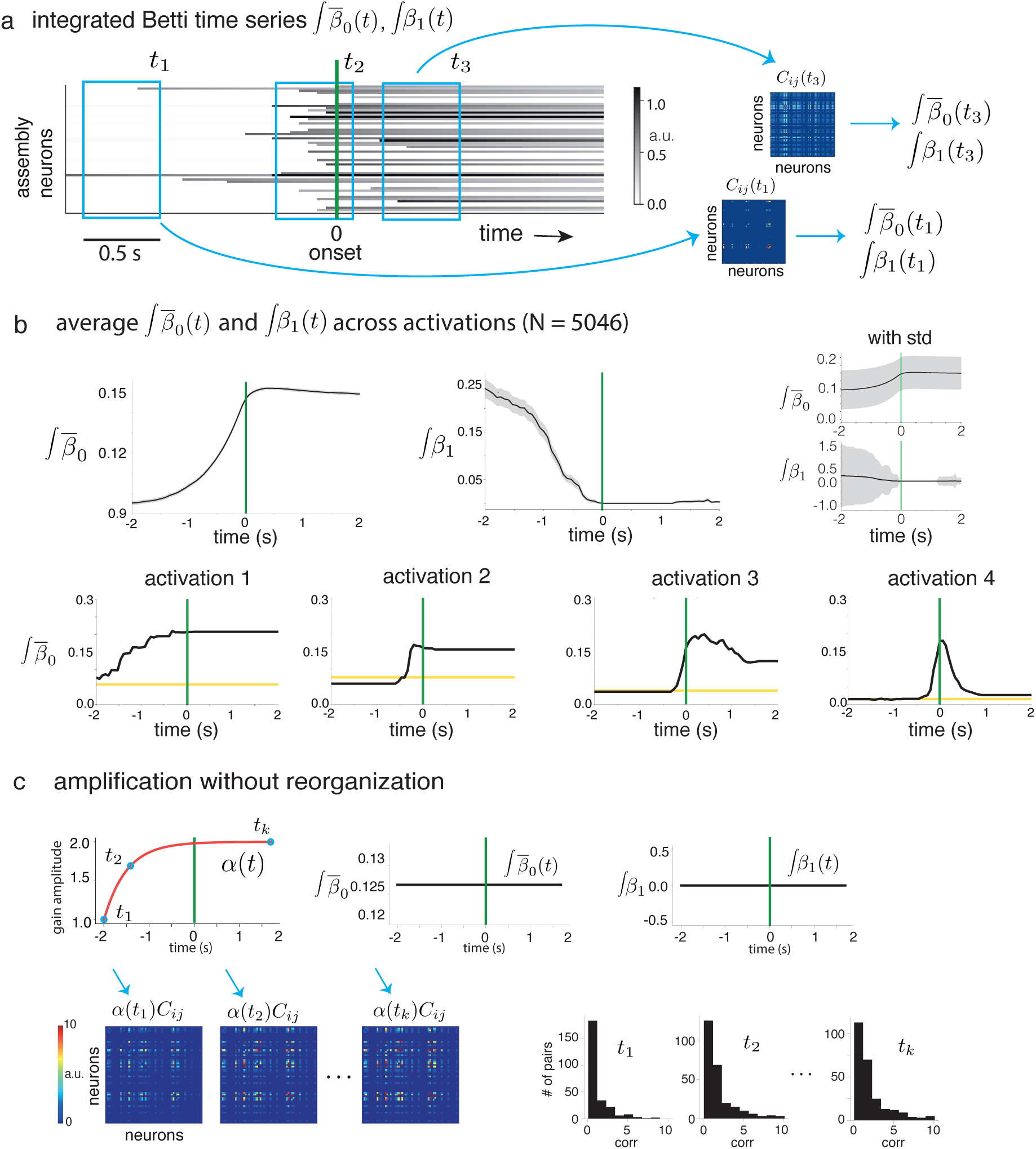
Topological dynamics of assemblies around spontaneous activations (a) Integrated Betti time series were computed from a sliding window correlation analysis. From each 0.5 s window we computed a correlation matrix *C_ij_*(*t*), where *t* is the end time of the window, and then from each *C_ij_*(*t*) we computed integrated Betti values 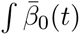 and *β*_1_(*t*), yielding integrated Betti time series. Activation onset is shown at time 0 (vertical green line). **(b)** (Top) The average 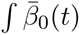 and *β*_1_(*t*) time series across assembly activations (N = 5046 for 12 larvae) standard errors shown in gray. While 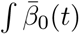 displays a sharp increase at activation onset, ∫ *β*_1_(*t*); shows a marked decrease to near zero. (Top right) The standard deviations (grey) of 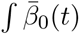 and∫ *β*_1_ (*t*) are shown together with the average curves (black). Note that ∫ *β*_1_ (*t*) goes from very high to very low variability following activation onset. (Bottom) Four examples displaying different dynamic profiles of 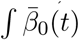 time series (black) around spontaneous activations. Yellow matrices of the same size as the activated assembly. **(c)** (Top) Amplification of correlations *C_ij_* over time via a multiplicative gain function *α*(*t*) = 1 − *e^βt^*(red curve) yields integrated Betti time series 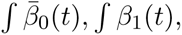 that are completely flat. (Bottom) Correlation matrices and distributions of correlation values at different time points (*t*_1_*, t*_2_*, t_k_*). Note that the amplification does not reorganize the structure of the correlation matrices, and thus the Betti curves remain unchanged.

The average integrated Betti time series 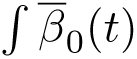 and *β*_1_(*t*) showed very different dynamics before and after activation onset (Figure 6b). 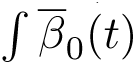 displayed a gradual increase until reaching a plateau at the onset of activation. In contrast, ∫ *β*_1_ decreased to almost zero just before activation onset. These trends were independent of the choice of sliding window size, and were consistent across larvae (Supp. Figure 9). The standard deviations prior to activation onset were large for both 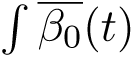 and *β*_1_(*t*), and remained large during activations for 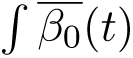 but decreased significantly for ∫ *β*_1_(*t*) (Figure 6b, top right). This suggests that spontaneous activations may display a variety of different dynamic activation patterns. Indeed, when we looked at the dynamic structure of individual activations, we observed different patterns of 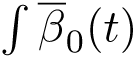 (Figure 6b, bottom).

It is worth emphasizing that the dynamics of 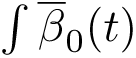 and ∫ *β*_1_(*t*) observed in Figure 6b reflected a reorganization of the assembly correlation structure around activation onset, not a strengthening of the correlation values. In fact, 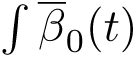 and *β*_1_(*t*) remained constant under amplification of the correlations without reorganization (Figure 6c).

To investigate whether the neuronal assemblies show a discrete number of potential dynamic structures during the spontaneous activations, we performed clustering of the 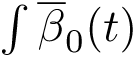 activation profiles (see Methods). For this analysis, we only used 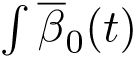 because ∫ *β*_1_ (*t*) displayed very little variability during activations. Using k-means clustering, we found that the 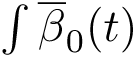 dynamics across all activations could be classified into six different groups (Figure 7a, see also Supp. Figure 10-11). The clusters we found had average 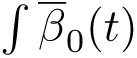 profiles that echoed the previously observed features of individual activations in Figure 6b. PCA analysis confirmed that the clusters grouped together activations based on features such as the rise (PC2) and fall (PC3) of 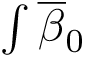 around the activation onset (Figure 7b). We observed that all six clusters can be represented in the activations of a single assembly; and, indeed, they all arose in each of the 12 larvae (Supp. Figure 12).

**Figure 7.**
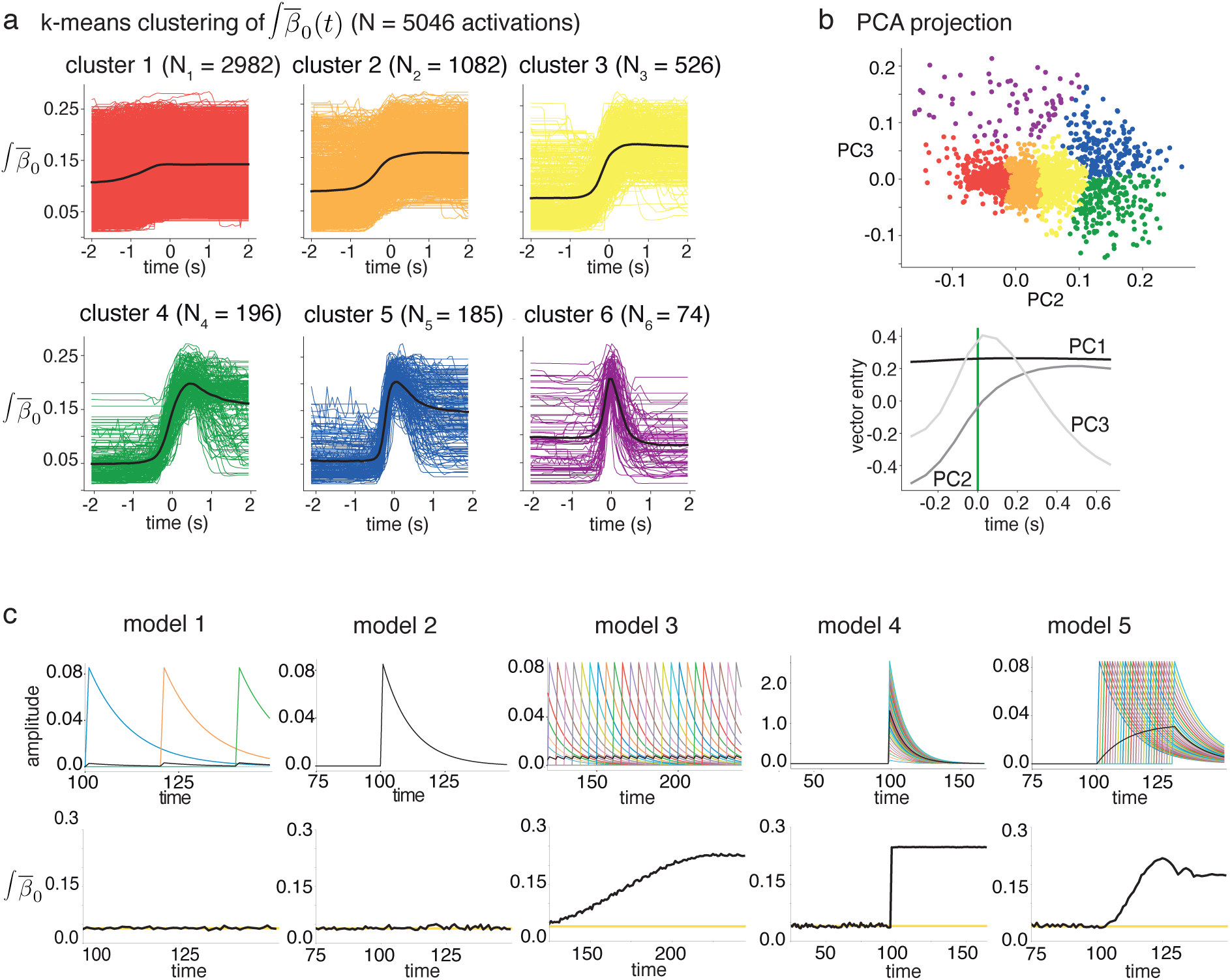
Clustering assembly activations by their. 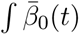 **time series. (a)** 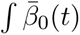 dynamic profiles (colors) for each cluster, together with their respective averages (black) (total N=5046 for 12 larvae). **(b)** (Top) PC2-PC3 projection of 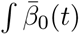 for all assembly activations (dots colored according to the clusters in (a)). (Bottom) Vector representations of PC1, PC2, and PC3. **(c)** (Top) A two-parameter family of toy models that vary in synchrony and amplitude of neuronal spiking. For each of five models, synthetic activity of 30 neurons and their average (black) is shown. (Bottom) The resulting 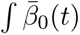 for each model (black). Yellow lines show expected 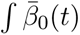 for a random symmetric i.i.d. matrix of size 30. Notice that a simple increase the in the average activity of the population does not correspond to an increase in 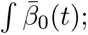 instead, a change in *β*_0_ reflects a change in the organization of correlations.

To learn about the mechanisms underlying the emergence of the different dynamic structures seen in the activation clusters (Figure 7a), we built a set of toy models of assembly dynamics where we varied the amplitudes of the neurons’ activity, and their synchrony (see Methods). We selected five models of varying amplitude and synchrony of neural spikes for assemblies of 30 neurons. Then, we calculated their 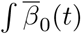 dynamics (Figure 7c). These models produced 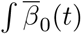 profiles similar to the variety observed across the tectal assembly activations (Figure 6b, Figure 7a). This suggests that the dynamic structure of the assembly activations can follow several activation patterns. For example, cluster 1 (Figure 7a, red) could be explained by a synchronous activation with neurons firing at similar amplitudes; cluster 2 (orange) could be explained by asynchronous activations while conserving the same amplitudes; cluster 3 (yellow) could be explained by synchronous activations with neurons firing at different amplitudes; and clusters 4 and 5 (green and blue) could be explained by rapid sequential firing of neurons at a conserved amplitude. However, cluster 6 (purple) did not have a dynamic profile matching any of the variants of the toy model.

To learn about the biological significance of the different types of activation dynamics, we computed for all six clusters the assembly sizes, activation lengths, and participation ratios (the fraction of active neurons in the assembly). We found that all clusters involved assemblies of similar sizes. However, clusters 4-6 showed longer activation lengths and larger assembly participation ratios (Figure 8a). We also found differences between clusters 1-3 versus 4-6 when considering the change in participation ratio between the periods prior to and after activation onset (‘pre’ and ‘post’, Figure 8b left). However, all clusters had similar E-I ratios prior to activation onsets, which were maintained during the activations (*P <* 0.1, ANOVA, p (pre): 0.224, p (post): 0.180, Figure 8b right).

**Figure 8.**
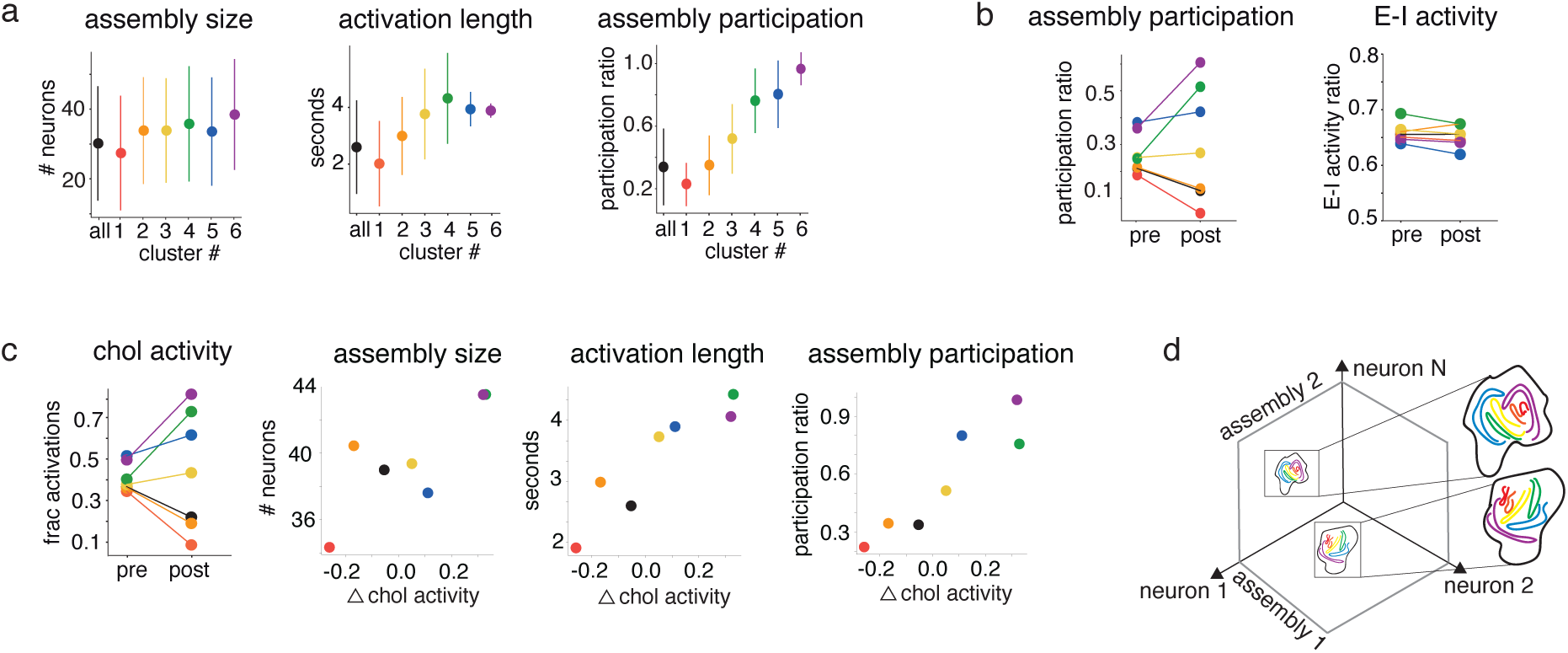
Cholinergic activity modulates assembly correlations resulting in longer activations with higher neuronal participation. **(a)** Average and standard deviation of assembly size, activation length, and participation ratio (fraction of active neurons of the assembly) in each of the assembly activation clusters from Figure 7a. **(b)** Average assembly participation and E-I activity ratio within the 2 seconds preceding (‘pre’) and following (‘post’) the onset of spontaneous activations. Note that clusters 4, 5, and 6 have a larger proportion of active assembly neurons after the activation onsets, yet the E-I balance is maintained from ‘pre’ to ‘post’ periods across all clusters. **(c)** (Left) Fraction of assembly activations containing active cholinergic neurons in the ‘pre’ and ‘post’ periods, for each cluster. Clusters 4, 5, and 6 show an increase in cholinergic engagement during the ‘post’ period. (Middle and Right) “Δ chol activity” denotes the change in the fraction of assembly activations containing active cholinergic neurons from assemblies that have at least one cholinergic neuron. While the change does not appear to correlate with average assembly sizes for the clusters, clusters with higher Δ chol activity have longer activation lengths and higher assembly participation. **(d)** The conceptual picture that emerges is of assemblies lying on low-dimensional subspaces, with each assembly exhibiting a variety of activation types belonging to distinct clusters, potentially modulated by cholinergic activity.

Next, we studied the effect of cholinergic activity on the properties of assembly activation clusters. Given the particular spatial distribution of the cholinergic neurons along the retinotopic tectal axis (Figure 1d), the presence of few cholinergic neurons within each of the assemblies (Figure 1b), and their strong but variable participation during spontaneous activations (Figure 2a and c), we hypothesized that cholinergic neurons could play a modulatory role in the dynamics of the tectal assemblies.

The fraction of activations per cluster involving cholinergic neurons diverged across clusters: despite being similar pre-onset, clusters 4-6 exhibited increased cholinergic activity post-onset while clusters 1-3 did not (Figure 8c, left; *P <* 0.01, ANOVA, P (pre): 0.041, P (post): 2.425e-99; see Methods). We also found that, across clusters, the change in cholinergic participation from pre-to post-activation onset (Δ chol activity) did not correlate with average assembly size (Figure 8c, second panel; Pearson corr: *r* = 0.84, p-value 0.0336). However, Δ chol activity did correlate with the average activation lengths and participation ratios (*r* = 0.93 and *r* = 0.98, respectively, *P <* 0.01; Figure 8c, right panels). This suggests that the participation of cholinergic neurons during the spontaneous activation of tectal assemblies modulates the number of recruited neurons and the duration of spontaneous activations (see also Supp. Figure 12). These results are consistent with a scenario where neural activity collapses onto lower-dimensional subspaces during assembly activations, yet within each assembly subspace the individual activations can exhibit a variety of distinct dynamic profiles (activation clusters) modulated by cholinergic activity (Figure 8d).

## Discussion

The tectal circuit is organized according to neuronal assemblies distributed along the circuit’s retinotopic map [28, 43, 29, 31]. The activations of tectal assemblies represent all-or-none preferred network states shaped by reciprocal inhibition. These characteristics promote an increase in strength and duration of visual responses, acting as a top-down fovea to improve visual resolution and prey-capture behaviors [31]. To learn about the structural connectivity principles underlying the functional role of these tectal assemblies, we identified the neurotransmitter identity of the neurons composing them (cholinergic, GABAergic and glutamatergic) and investigated their activity using a topological data analysis approach.

We found that assemblies have a higher E-I ratio as compared to the entire tectal circuit, indicating that assemblies correspond to highly excitable subcircuits with inhibition serving mainly to counterbalance excitation, as in a balanced state [48, 49, 50, 51, 52]. The cholinergic neurons were distributed along the tectal retinotopic axis, resulting in a majority of tectal assemblies containing at most a handful of cholinergic neurons [53, 40]. Despite a stable E-I ratio, we found that the structure of neuronal correlations undergoes a rapid and dynamic reorganization near the onset of assembly activations. This was evident from our topological analyses and the rise in entropy of the assembly activity patterns at the start of activations.

For the topological analyses of functional connectivity, we used Betti curves to quantify the topological features of pairwise correlations of tectal assemblies. Like other methods from topological data analysis, Betti curves can be used to uncover intrinsic higher-order geometric and topological organization of neural activity and are remarkably robust to noise and the underlying nonlinearities of the data [36, 46, 34]. Comparison between the Betti curve results for the assemblies with those of various synthetic models of different ranks showed that assembly corre-lations during activations (‘on’ periods) are consistent with a low-rank functional connectivity structure [47]. On the other hand, outside of activations (‘off’ periods) the assembly correlations more closely resembled those of randomized assemblies and were more consistent with high-dimensional comparison models. Notably, the underlying low-rank structure we detected via topological methods is not visible with traditional spectral methods because of nonlinearities in the calcium imaging data [47] (Supplementary Figure 7).

Several studies have suggested that low-rank connectivity matrices provide a mechanism for generating low-dimensional dynamics, and that this in turn can lead to attractor-like dynamics [54, 55]. More generally, low-dimensional dynamics have been reported in a variety of neural systems, particularly when the animal is engaged in a specific task [56, 57]. The picture that emerges from our work, however, is not that of a network with a single low-dimensional manifold of neural activity. Instead, each assembly activation moves the tectal network activity towards a distinct low-dimensional subspace corresponding to the assembly. The reduction in dimensionality is two-fold: first, there is a reduction from the tectal circuit to the assembly (∼20-90 dimensions as opposed to thousands, representing the spatial location of the visual stimulus); second, there is a further reduction to the low-rank structure of the pairwise correlations within each tectal assembly. In other words, the population activity is high-dimensional, but individual assembly activations (whether spontaneous or stimulus-driven) push the network into distinct low-dimensional dynamic regimes (Figure 8d).

To investigate the dynamics of the topology of assembly activations in more detail, we implemented a novel TDA approach using integrated Betti time series, 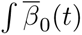 and ∫ *β*_1_(*t*), which track the temporal changes in assembly correlations around activation onsets (Figure 7). First, we found that both the 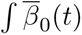 and ∫ *β*_1_(*t*) time series were highly stereotyped across all activations, and that activations could be clustered into six distinct groups using the 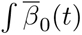 time series. In a toy model, we observed that the different dynamic profiles for 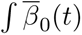 could be recreated by modifying the temporal patterns and amplitudes of the activations. This suggests that different clusters may represent different functional roles or coding capacities of the assemblies. We found that some of the 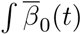 activation profiles displayed longer durations and a larger proportion of active neurons during spontaneous activations. Interestingly, these profiles were also associated with the activation of cholinergic neurons within the assemblies (Figure 8). Previous studies have shown that attention is associated with the activation of the cholinergic system [58, 59]. Our results suggest a mechanism by which attention may influence visual detection via the modulation of the topological structure of sensory circuits.

## Acknowledgments

We thank D. Souchet for help with zebrafish; M. Privat and E. Marachlian for help with the light-sheet microscope and discussions; and J. Londoño Alvarez, C. Lienkaemper, and J. Paik for discussions. We also thank the IBENS imaging facility. This work was supported by the US-France CRCNS grant NIH R01 NS120581 (to CC) and ANR-20-NEUC-0001 (to GS). CC was additionally supported by NSF DMS-1951165 and a Simons Fellowship, and GS was supported by ERC CoG 726280.

## Methods

### Experimental methods – calcium imaging

All experiments were performed using transgenic zebrafish larvae from 6 to 7 days post-fertilization (dpf). The embryos were collected and raised at 28^◦^C in 0.5x E3 embryo medium (E3 in mM: 5 NaCl, 0.17 KCl, 0.33 CaCl_2_, 0.33 MgCl_2_, pH 7.2). Larvae were kept under a 14/10 h light-dark cycle and fed after 5 dpf with paramecia. Zebrafish sex cannot be determined until 3 weeks dpf. Thus, the sex of the animals is unknown. All experimental procedures were approved by the Comité d’éthique en expérimentation animale *n*^◦^ 005. Reference number APAFIS#27495-2020100614519712 v14.2.2.

### Transgenic lines

We used double transgenic (HuC:H2B-GCaMP6f and Vglut2a:loxP-mCherry-loxP-Gal4) zebrafish larvae in a nacre background (gift from the National BioResource Project, Zebrafish, Core Institution).

### Selective-plane illumination microscopy for calcium imaging

We recorded the sponta-neous activity of the zebrafish optic tectum using selective-plane illumination (SPIM) microscopy with near single-cell resolution [60, 61]. The calcium indicator (H2B-GCaMP6f) was excited by a micrometer-thick light sheet emitted from the left side of the larva which produced optical sectioning.

We used a constant wave 488 nm laser (Omicron PHoXX® 480-200) as the single-photon light source of the SPIM. The laser beam was coupled to an optical single-mode fiber and the output was collimated using a fiber optic. The 1.6 mm beam from the coupled laser was expanded to a 4.5 mm beam through a telescope formed by two lenses (*f*_1_ AC254-050-A-ML and *f*_2_ AC254-150-A-ML, Thorlabs). The telescope created a collimated beam with uniform intensity.

The laser beam was projected onto a pair of galvanometric mirrors at an angular range of 26^◦^ (Model 6215H with a mirror size of 5 mm). A system composed of a scan lens (AC508-080-AB-ML, Thorlabs) and a tube lens (Olympus U-TLU IR) imaged the beam to a decentered spot at the back focal aperture of the imaging objective.

The scan lens (*f*_scan_ 50.8 mm) was placed in front of the scanning mirror and converted the angular deflection *θ* into a horizontal line displacement of the incident light. The beam was collimated by an infinity corrected tube lens (*f*_tube_) and steered onto the back focal plane of the objective lens.

The generated fluorescence of the sample was collected by a Hamamatsu ORCA-Flash4.0 V2® camera whose optical axis was orthogonal to the excitation plane. We used the *µ*Manager software to control the Orca-Flash4.0 camera.

### Recording

We recorded at 15 Hz the spontaneous brain activity of the optic tectum for one hour in zebrafish larvae aged between 6 to 7 dpf. The zebrafish larvae were embedded in 2% low-melting point agarose (Invitrogen) in E3 embryo medium [62] (in mM): 5 NaCl, 0.17 KCl, 0.33 CaCl_2_, 0.33 MgCl_2_ (pH 7.2). The larvae were placed inside a rectangular chamber filled with E3 embryo medium. This chamber was placed on an elevated stage. We recorded a volume of 16 µm of the optic tectum starting, on average, 52.8 µm (±15.26 µm SD) below its surface. The volume consisted of 3 layers separated by 8 µm. Larvae were left to rest and adapt to the recording environment for at least 30 min before starting the experiments.

### Registration of recorded images

We used a template based alignment [63] to register the raw images. While most of the images were aligned to the template, some images (in particular those recorded during movements and display movement artifacts) had a low correlation with the template. For that reason, we z-scored all of the correlation values obtained in the previous step. Then we looked at the distribution and set a threshold. All images whose correlation values were below that threshold were removed. Finally, we inspected the remaining images to identify and remove other images that could be attributed to artifacts.

### Experimental methods – neurotransmitter identity

The optic tectum contains neurons of three different types of neurotransmitter identity: cholinergic, GABAergic, and glutamatergic [53, 40]. The glutamatergic neurons were identified *in vivo*, while the cholinergic and GABAergic neurons were identified via immunostaining.

### Detection of Vglut2a neurons

We identified Vglut2a neurons based on the expression of the red fluorescence protein mCherry. Using the SPIM, we imaged a volume of the optic tectum of 48 µm depth which comprised 13 layers that were separated by 4 µm.

We matched the GCaMP6f images with the mCherry images to select the proper layers. Then we split the mCherry images into different regions to take into account differences between intensities. We calculated the mean and standard deviation of each region and then fixed a threshold to detect the brighter areas. Then, we calculated the mean intensity of the detected neurons inside each area. Finally, those neurons whose mean intensity was equal or higher than the threshold of their area were labeled as glutamatergic neurons. The threshold was the mean plus 1/2 of the standard deviation of the detected neurons for each region. Later, we used the immunostaining results to double check that the detected neurons were glutamatergic.

### Immunostaining

Ten double transgenic (HuC:H2B-GCaMP6f and Vglut2a:loxP-mCherry-loxP-Gal4) zebrafish larvae were anesthetized and fixed in a solution of PBS+4% PFA at a pH of 7.4. Next, the fixed larvae underwent a series of washes in PBS+0.5% Triton X-100 to remove the fixative. A blocking solution containing PBS+0.5% Triton X-100, 1% DMSO, and 2% BSA was applied to saturate nonspecific binding sites. The samples were then incubated with primary antibodies diluted at 1/100 against specific markers of interest, such as Rabbit Anti-Gad1b, Goat Anti-ChAT, and Mouse Anti-mCherry. After that, the samples were washed again and subjected to a secondary antibody staining step using Donkey Anti-rabbit 647, Donkey Anti-goat 568, and Donkey Anti-mouse 488 antibodies diluted at 1/500. After further washes, the samples were treated overnight with 500 µL of PBS, 1% Triton, 1% DMSO, 0.1% Tween, and DAPI (10 µg/µL) for nuclear staining. Finally, the samples were fixed once more with PBS+4% PFA, washed, and stored in PBS-0.5% solution at 4^◦^C until confocal imaging.

The samples were immobilized in 2% UltraPure™ Low Melting Point Agarose (16520100, Invitrogen™) on a glass-bottom µ-Dish 35 mm (high Glass Bottom) (81158, ibidi). Imaging was conducted using an inverted Confocal Laser Scanning Microscope TCS SP8 (Leica, Germany), equipped with a CS2 Plan Apochromat 40x/1.10NA water immersion objective lens and utilizing hybrid PMTs. To minimize any potential crosstalk between channels, images were acquired in frame sequential mode. All images were captured using a White Light Laser (WLL) with excitation wavelengths of 488 nm, 568 nm, and 647 nm to excite Alexa Fluor Plus 488, Alexa Fluor 568, Alexa Fluor Plus 647, respectively, as well as a 405 nm diode for DAPI excitation.

The pinhole was set at 1.0 Airy unit. Imaging was performed with a 2 µm Z-step interval at a resolution of 1024 × 1024 pixels. The zoom factor was 0.75. The immunostaining images were analyzed using the image processing package Fiji [64]. We aligned the images using the Fiji plugin BigWarp [65]. Neurons that were not identified as ChAT or Vglut2a positive were considered as GABAergic.

### Data analysis methods – preprocessing

#### Segmentation

To segment single neurons, we used the algorithm introduced in [66] using the default parameters except for the value of the initial radius, which we set to 2 um instead of 3.1 um. Briefly, we used the template for registration to detect GCaMP-positive areas. These areas were coarse-extracted by binary thresholding based on absolute pixel intensity and local image contrast. Then, local intensity normalization was performed on each pixel by calculating a relative rank of pixel intensity. The normalized image was smoothed by an averaging kernel and a pixel with a maximum value within a surrounding image patch was identified as a centroid of a neuron. This local smoothing and maxima was repeated twice to maximize the number of neurons correctly recognized. After detecting neurons, we imported the neurons to the Toolbox described in [41, 42] where time series of pixel intensity were averaged from pixels within each individual neuron. This gives the fluorescence time series of each neuron.

#### Calculation of ΔF/F

To calculate the ΔF/F for each fluorescence time series, we first calculated the baseline neural fluorescence by computing a running average of 2700 time points (∼ 180 seconds) for the 8*^th^*percentile of the data [67]. We then calculated the change in fluorescence relative to baseline fluorescence, ΔF, divided by baseline fluorescence, F, to get ΔF/F. Due to the preprocessing of the fluorescence traces, the estimated baseline fluorescence can be very small or negative in rare cases. To prevent spurious ΔF/F estimates, we set the baseline as the maximum of the median filter estimated baseline and the standard deviation of the estimated noise of the fluorescence.

#### Detection of calcium transients and deconvolved calcium signal

We applied the algorithm described in [68] to the ΔF/F to detect the calcium transients (events). Neurons without calcium transients were removed from the analyses. In addition to the calcium transients, the algorithm calculates the estimated calcium concentration (deconvolved calcium) using the ΔF/F signal. The deconvolved calcium signal is a denoised and smoother version of ΔF/F; we used it to compute neuronal correlations for topological analyses.

#### Assembly detection

We detect neuronal assemblies by following the PCA-PROMAX approach described in [28, 42, 29]. First, we reduced the dimensionality of the neuronal ΔF/F dynamics by using principal component analysis (PCA) and then only keeping the principal components with a statistically significant eigenvalue. We then softened the PCA orthogonality condition by rotating the principal components using PROMAX. Finally, we set a data-driven threshold on the PCA-PROMAX loadings to establish the neuronal compositions of the assemblies. We used the Computational Toolbox introduced in [28, 41, 42]. (See https://github.com/zebrain-lab/Toolbox-Romano-et-al.)

#### Compact assemblies

To distinguish between sparse and compact assemblies, for each assembly we computed the distribution *P* of pairwise Euclidean distances between the neurons. We then computed the mean and Shannon entropy, 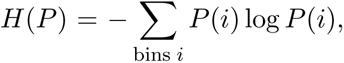 of each (binned) distance distribution, obtaining a pair of values (entropy, mean distance) for each assembly.

Next, we examined the joint distribution of these values across all identified assemblies (see Supp Figure 1e). This distribution was bimodal, corresponding to distinct collections of compact (purple) and sparse (green) neuronal assemblies. We found by visual inspection that assemblies with an entropy value lower than 1.5 were spatially concentrated (*compact assemblies*). The compact assemblies had distance distributions with both lower entropy and lower mean distance compared to the other (sparse) assemblies.

#### Detection of assembly activations

We detected spontaneous activations of assemblies using criteria combining the percentage of active neurons in the assembly with the matching index (MI), a measure of how much of the total population activity is coming from a given assembly.

For an assembly *i* and time point *t*, the MI quantifies the overlap between the binary activity pattern of the assembly, Pat*_i_*(*t*), and that of the entire recorded population, Pat_pop_(*t*), where Pat*_i_*(*t*) and Pat_pop_(*t*) are index sets of active neurons, and |Pat*_i_*(*t*)| denotes the size of the set. Specifically, the MI is defined as

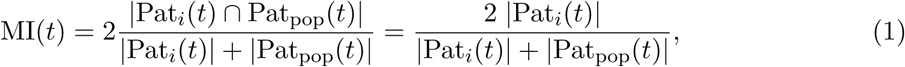

where the second equality follows because Pat*_i_*(*t*) ⊆ Pat_pop_(*t*). Note that MI(*t*) always lies in [0, 1], with 0 indicating no overlap and 1 indicating complete overlap (Pat*_i_*(*t*) = Pat_pop_(*t*)); we can thus refer to the value as a percentage.

We defined the activation *onset* of an assembly as the time when the percentage of active neurons in the assembly reaches at least 15% and the value of the MI is at least 10% [28]. The activation *offset* was defined as the time at which the percentage of active neurons in the assembly decreases below 12.5% or the MI value drops below 10%. A neuron is deemed active for 3 seconds after the detected calcium events. We do not consider activations in our analysis that have time frames with movement artifacts.

For each assembly, the ‘on’ and ‘off’ periods are defined as the time periods during or between activations, respectively (see Figure 5a).

#### Assembly and activation selection

10 of the 12 larvae were successfully immunostained, so that we were able to distinguish all three cell types: glutamatergic, GABAergic, and cholinergic.

For the analyses in Figures 1-2, we used the 10 successfully immunostained larvae and focused on assemblies with at least 10 neurons (458 of the 517 assemblies, see Supp. Figure 1b).

For the analyses in Figures 4-8, we used all 12 larvae, including the two where immunostaining was unsuccessful. In Figures 4-5, we restricted our analyses to assemblies with at least 20 neurons (N = 450 assemblies across the 12 larvae), as Betti curves with just a few neurons are difficult to compare to those from larger assemblies (see Supp. Figure 4).

In Figures 6-8, we considered all spontaneous activations, including those from assemblies with fewer than 20 neurons (N = 5046 activations across 450 compact assemblies for 12 larvae).

### Data analysis methods – E-I balance, entropy, correlations

#### E-I analysis

We analyzed the excitatory-inhibitory (E-I) ratio, using the formula *E/*(*I* + *E*), from three seconds prior the onset of activations to ten seconds after the onset using the ΔF/F signal. We divided the time period in non-overlapping windows of one second each. We assessed whether the difference between the medians of the windows was statistically significant by using Kruskal-Wallis test (a non-parametric version of ANOVA that does not assume normality). To identify which windows differed significantly from each other, we used the Dunn test post-hoc (Holm correction) in Figure 2d.

#### Entropies of assemblies during activations

For a single activation from a given assembly, we computed entropy time series as follows. First, we used a causal sliding window of 0.5 s, with slide step of one time bin (≈ 67 ms), and summed the activity of each neuron to compute a population vector for each time window. Normalizing by the total population activity, this yields a distribution *p* = (*p*_1_*,……, p_n_*) for each time window describing the fraction of total activity coming from each neuron. For each such distribution, we computed the Shannon entropy 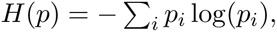 which satisfies

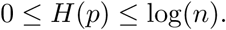

Because neuronal assemblies are composed of different numbers of neurons, we normalized the entropy by the maximum value, log(*n*), to obtain:

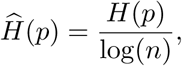

with 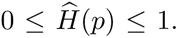. Collecting the entropies across time windows, we obtain an entropy time series for each activation that can be averaged across activations (Figure 2e). We similarly computed entropies for subpopulations corresponding to distinct neuron types (glutamatergic, GABAergic, cholinergic). We did not include the cholinergic populations when the assembly had only one cholinergic neuron. In addition, we did not take into account the entropy of the cholinergic population after 4 s of the activation onset because of the reduced number of samples.

#### Correlations

In Figure 1c we used Pearson correlation. For the topological analyses, we computed the pairwise correlation between neurons *i* and *j* as

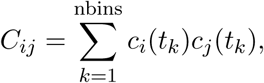

where *c_i_*(*t*) is the deconvolved calcium signal of neuron *i* at time *t*. We did not mean-shift *c_i_*(*t*), as is done in Pearson correlation, because the neural activity is sparse and mean shifting would result in co-silent neurons having high correlation. Since *c_i_*(*t*) decays slowly, we also did not require further smoothing of the signal. In particular, integrating cross-correlograms over a 1 second window of delays as in [36] did not yield a noticeable difference in the results of our topological analyses. All correlations were computed after removing time bins with movement artifacts.

Assembly correlations for ‘on’ and ‘off’ periods, as in Figure 5, were computed by restricting the above *C_ij_*computations to time bins during (‘on’) or between (‘off’) activations. Since there are far fewer ‘on’ than ‘off’ time bins for any given assembly, the ‘off’ correlations were computed by randomly subsampling the time bins between activations so that their total matched the number of ‘on’ time bins.

#### Randomized assemblies

For each compact assembly of size *k*, as a control we considered *randomized assemblies* that were constructed by taking a random sample of *k* neurons from the full population of neurons (we only sampled from neurons that were active during the hour-long recording). We also considered random samples of neurons that contained the same numbers of glutamatergic and non-glutamatergic neurons as the compact assemblies, and observed that this did not significantly change the resulting analyses.

#### Correlation shuffles

As another control, for each compact assembly we randomly shuffled the matrix of correlations (entrywise) in the upper triangle, [*C_ij_*]*_i<j_*, to obtain a symmetric shuffled correlation matrix with the same distribution of correlation values as the original assembly. In particular, any analysis of correlations that depends only on the distribution of values would produce identical outputs for the original and shuffled correlation matrices (see Figure 3d).

#### Low-rank and high-rank/dimension matrix models

We generated random *n* × *n* correlation matrices *A* of rank or underlying dimension *d* by selecting *n* points 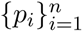 uniformly at random from either [−1, 1]*^d^* (‘mixed’ model) or [0, 1]*^d^* (‘positive’ model) and setting *A_ij_* = ⟨*p_i_, p_j_*⟩ where 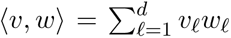 is the standard inner product on 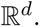 Note that the ‘mixed’ model is the standard procedure for generating random matrices of a fixed rank, while the ‘positive’ model more closely aligns with correlation matrices arising from calcium imaging data, as the signal has nonnegative values. In Figure 5c-d we considered low ranks *d* = 1*,…,* 5; medium ranks *d* = 20 − 90 that match the distribution of assembly sizes; and high underlying dimensions of *d* = 500 and *d* = 54, 000, where 500 is roughly the number of time bins of an assembly activation and 54, 000 is the number of time bins in the full recording. Note that while the maximum rank of an *n* × *n* matrix is *n*, the Betti curves distinguish models whose underlying dimensions are considerably higher [36].

#### Software

We carried out all analyses in this research by using a combination of MATLAB and Python scripts. We used some scripts of the Calcium Toolbox (described in [41, 42]) for the processing of the fluorescence data. The segmentation of the neurons was done with the method introduced in [66].

### Topological data analysis methods

#### TDA software

We computed Betti curves *β_i_*(*ρ*) for symmetric correlation matrices following the clique topology pipeline in [36, 34] using our Python package, PyCliqueTop 2023, which is freely available at: https://github.com/nerdnik/PyCliqueTop_2023. This code is an updated Python version of the MATLAB package CliqueTop, first introduced in [36] and used in [69, 70]. Whereas CliqueTop uses Perseus to compute persistent homology, PyCliqueTop 2023 uses the Python package Ripser for faster, more memory-efficient computations.

#### Averaging of Betti curves

The Betti curves *β_i_*(*ρ*) coming out of the software are discretized vectors that partition the [0, 1] interval of possible edge densities in different ways. If *n* is the number of neurons in an assembly, then 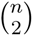 determines the discrete set of edge densities for each step of the filtration. To compute average Betti curves across assemblies with different numbers of neurons, we linearly interpolated the discretized vectors to get Betti curve values for evenly spaced edge densities 0 to 1 with step size 0.001.

#### Integrated Betti values

To obtain a single summary statistic from each Betti curve, *β_j_*(*ρ*), we used the integrated Betti value:

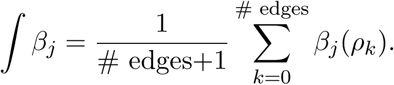

These values were also used in [36, 69, 70] to simplify the statistics when comparing Betti curves to null models.

To fairly compare *β*_0_ curves and ∫ *β*_0_ values across assemblies of different sizes, we first normalized the *β*_0_ curve by dividing by the size of the assembly (# neurons), and then subtracted 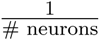 from the integrated *β*_0_ curve. We thus obtained:

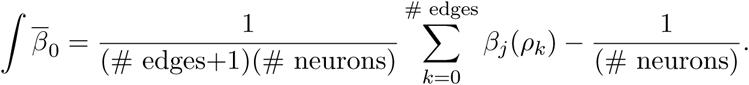

#### Integrated Betti time series clusters

This is a novel TDA analysis that has not been described elsewhere. For each activation, we computed correlations in a 0.5 second causal sliding window starting 2 seconds prior to activation onset through 2 seconds after, with a slide step of 1 time bin (≈ 67 msec). The time windows are indexed by the final time bin, *t_i_*. As we recorded at 15 Hz, this results in 61 temporally ordered assembly correlation matrices, *C_ij_*(*t_i_*), per activation (see Figure 6a). We checked the effects of the sliding window size on the results of our analysis, considering window sizes of 1 time bin, 4 time bins (≈ 0.25 s), 8 time bins, 15 time bins (1 s), and 30 time bins (see Supp. Figure 9).

The correlation matrices *C_ij_*(*t*) for a given activation were run through the clique topology pipeline, and we restricted attention to the two summary values 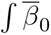 and ∫ *β*_1_ for e_∫_ach matrix.

Pooling together these values across *t* results in two time series per activation, 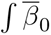 ∫ *β*_1_(*t*).

We then clustered the activations via their 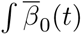 time series using k-means clustering, viewing each time series as a vector (Figure 7a). Before clustering, we first restricted each 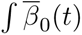 to a small window starting 5 time bins (≈ 0.33 s) before activation onset to 10 time bins (≈ 0.67 s) after activation onset. We then subtracted the mean and clustered the resulting vectors using k-means clustering. To determine the number of clusters, we ran a silhouette analysis (see Supp. Figure 10). We also checked the effects of the choice of restriction window on the k-means clusters (see Supp. Figure 11). Additionally, we performed PCA on the restricted 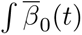 time series (Figure 7b). However, we did not mean-center as we did for the k-means clustering. We observed that the first principal component corresponded to the mean of confirming the importance of mean-centering 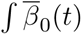 prior to k-means clustering.

#### Synthetic models for integrated Betti time series analysis

We compared the integrated Betti time series for assembly activations to those of synthetic activation dynamics generated by a family of simple spiking models with varying degrees of synchronicity and amplitude. We generated synthetic calcium signals from a set of *N* = 30 spike trains where we varied the synchrony and amplitude of the spikes. We set a sampling frequency of 15 Hz and convolved each spike train with a causal decaying exponential with timescale *τ* = 0.75 seconds over a convolution window of *T* = 10*τ* seconds. We considered 5 models: model *k* has spike trains where neuron *i* spikes at time bin 100 + *λ_k_i* with amplitude 1 + *γ_k_i*. The model 1 parameters were (*λ*_1_*, γ*_1_) = (20, 0); model 2 had (*λ*_2_*, γ*_2_) = (0, 0); model 3 had (*λ*_3_*, γ*_3_) = (5, 0); model 4 had (*λ*_4_*, γ*_4_) = (0, 1); and model 5 had (*λ*_5_*, γ*_5_) = (1, 0) (see Figure 7c). For each of these model activations, we computed the 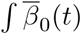 time series as described above.

#### Cholinergic activity by cluster

To compute the fraction of activations with cholinergic activity in ‘pre’ or ‘post’ activation periods (see Figure 8c), we computed the number of activations within each cluster that had at least one cholinergic assembly neuron active within two seconds prior to onset (‘pre’), and during activation (‘post’), and then divided each number by the number of cluster activations coming from assemblies that have at least one cholinergic neuron.

## Notes

### Competing Interest Statement

The authors have declared no competing interest.

