## Supplementary Figures for "Topological analysis of neuronal assemblies reveals low-rank structure modulated by cholinergic activity"

### Supplementary figure legends

**Supplementary Figure 1. Properties of neuronal assemblies.** **(a)** Optical section of a 6 dpf larva immunostained showing cholinergic neurons in the optic tectum. Yellow arrows point to eight cholinergic neurons. **(b)** (Top) Histogram of the total number of assemblies for ten larvae. The mean number of assemblies was 51.7 and the standard deviation 14.35. (Bottom) Histogram of the number of neurons in assemblies. We looked at the 517 assemblies of the 10 larvae and we found that an average assembly had $31.53 \pm 17.85$ neurons (mean $\pm$ SD). **(c)** Histograms of each cell-type proportion in assemblies. Black vertical lines represent the proportion of cholinergic, GABAergic, and glutamatergic neurons in the assemblies. Yellow vertical lines and shading represent the mean $\pm$ standard deviation of the proportion (across larvae) of each cell type in the periventricular layer. On average, the proportion of glutamatergic neurons is higher in the assemblies than in the periventricular layer. Conversely, the proportion of GABAergic neurons in the assemblies is lower than in the population. **(d)** Boxplots of the proportion of glutamatergic neurons in the assemblies for each larva. Dashed horizontal line represents the mean proportion of glutamatergic neurons in the periventricular layer (across larvae). In all larvae, the proportion of glutamatergic neurons in the assemblies was higher than in the periventricular layer. **(e)** To define the compact assemblies, we analyzed the distributions of pairwise distances between neurons within assemblies and found that entropy values of these distributions were correlated to mean distances, and could thus be used to separate the compact versus sparse assemblies. Each dot in the scatterplot represents the pair (entropy, mean distance) for one assembly. An entropy cutoff of 1.5 was chosen to define the compact assemblies (purple, entropy $\leq$ 1.5). Assemblies below this cutoff were verified as compact via visual inspection. **(f)** Graph analysis of differences between Pearson correlation matrices computed from $\Delta$F/F for the 'on' and 'off' periods of compact assemblies. Graphs were generated by including edges for high pairwise correlations up to an edge density threshold ($\rho = 0.75, 0.5, 0.25$). Erdös-Rényi random graphs with matching edge probabilities were generated as controls. (Right) Histograms show the differences in the numbers of communities identified for graphs from 'on' and 'off' assembly correlations versus control graphs. Communities were detected using the *Infomap* method [1]; the Python version of *igraph* was used for building the graphs and calculating graph measurements [2].

**Supplementary Figure 2. Betti curves reflect subcluster structure.** **(a)** Synthetic time series $c_i(t)$ representing the calcium signal for six neurons display two groups of similarly behaving neurons: neurons 1-4 are all co-active while neurons 5 and 6 are relatively silent, and vice versa. **(b)** Correlation matrix $C_{ij} = \langle c_i(t), c_j(t) \rangle$. Note that we set the diagonal to white as those entries are not used in the topological analysis. **(c)** The filtration of clique complexes associated to the correlation matrix $C$. At each step of the filtration, a single new edge (magenta) and any resulting cliques, including triangles and tetrahedra, are added to the simplicial complex (only edges are shown). Since all information about the clique complex is contained in the graph, we do not shade in these higher-dimensional simplices. Notice that the clique complexes in the filtration from $\rho = 4/15$ through $\rho = 7/15$ (circled in green) have exactly two connected components, one consisting of neurons $\{5, 6\}$ and the other with neurons $\{1, 2, 3, 4\}$, corresponding to the two groups of co-active neurons. These two connected components arise in the filtration because of higher correlations within these two clusters than across. At edge density $\rho = 8/15$, we have one connected component (circled in brown). **(d)** Betti curves $\beta_0$ (grey), $\beta_1$ (blue), and $\beta_2$ (red) for the filtration of clique complexes in (c). Observe that $\beta_0$ plateaus at a height of 2 from $\rho = 4/15$ through $\rho = 7/15$ (green vertical lines), dropping to 1 at $\rho = 8/15$ (vertical brown line), indicating that there are two 0-cycles, or connected components, for this period of the filtration. In this sense, plateaus in $\beta_0$ capture sub-cluster structure in correlation matrices.

**Supplementary Figure 3. Betti curves reflect periodic sequential firing.** **(a)** Synthetic time series $c_i(t)$ representing the calcium signal for six neurons with periodic sequential firing. **(b)** Correlation matrix $C_{ij} = \langle c_i(t), c_j(t) \rangle$. **(c)** The filtration of clique complexes associated to the correlation matrix $C$. A 1-cycle (cyan) emerges at $\rho = 6/15$ and disappears at $\rho = 11/15$ (circled in green). Note that homology cycles are actually equivalence classes of cycles, so there is not necessarily a unique representative set of edges that we can associate with a 1-cycle. For example, at $\rho = 0.67 = 10/15$ the cycles 1234 and 2356 are equivalent, and only count as one homology 1-cycle towards $\beta_1$. The added edge from 3 to 6 at $\rho = 11/15$ can be thought of as either "coning off" the 1234 1-cycle, or "filling in" the 2356 1-cycle via two triangles. **(d)** Betti curves for the filtration of clique complexes in (c). Notice that $\beta_1$, which tracks the number of homology 1-cycles in the filtration, begins at zero and increases to one from $\rho = 0.4 = 6/15$ to $\rho = 0.73 = 11/15$, and then returns to zero. This long-lasting 1-cycle reflects the sequential periodic firing within the neural population.

**Supplementary Figure 4. Average Betti curves for assemblies across the whole recording - per larva.** (Top) Average Betti curves, $\bar{\beta}_0$ (black), $\beta_1$ (blue), and $\beta_2$ (red) for assembly correlations versus randomized assembly correlations across all 12 larvae, reproduced from Figure 4b for comparison. (Bottom) Average Betti curves for assemblies and randomized assemblies for each larva. The per larva results are consistent with the overall averages.

**Supplementary Figure 5. Assembly size effects on integrated Betti values.** To check how the size of assemblies affects the integrated Betti values of assembly correlations (left) and randomized assembly correlations (middle), we show scatter plots of the assembly size versus $\int \beta_0$ (top) and $\int \beta_1$ (bottom). We observe moderate correlation between assembly size and $\int \beta_0$ for both assemblies ($R^2 = 0.31$) and randomized assemblies ($R^2 = 0.41$), and no correlation between $\int \beta_1$ for either ($R^2 = 0.01$ and $R^2 = 0.00$, respectively). Therefore we choose to use a normalized integrated Betti 0 value, $\int \bar{\beta}_0$, that we get by dividing $\int \beta_0$ by the size of the assembly and then subtracting $\frac{1}{\text{assembly size}}$. While size effects still remain, the correlation between assembly size and $\int \bar{\beta}_0$ is reduced for both assemblies and randomized assemblies ($R^2 = 0.22$ and $R^2 = 0.23$, respectively). Notice that these trends are not observed for the entrywise correlation shuffle of the assembly correlations (right), as there is no correlation between assembly size and $\int \beta_0$ ($R^2 = 0.00$) yet strong correlation between assembly size and $\int \beta_1$ ($R^2 = 0.96$). Instead, for the correlation shuffle random control, normalizing $\int \beta_0$ introduces a strong correlation between assembly size and $\int \bar{\beta}_0$ ($R^2 = 0.83$).

**Supplementary Figure 6. Sensitivity of TDA statistics to matrix size and robustness of low rank observations to choice of metric. (a)** To check how the assembly size (matrix size) affects integrated Betti values of low rank random matrix models, we show box-plots of the distribution of $\int \bar{\beta}_0$ (top), $\int \beta_1$ (middle), and $\int \beta_2$ (bottom) for matrices of sizes $25, 50$ and 75 for ranks 1-5 (columns) for each the 'mixed' and 'positive' models (100 matrices for each box plot). In the 'mixed' models, we observe a downward shift in the distribution of $\int \bar{\beta}_0$ with an increase in matrix size for all ranks. The rank 1 'positive' model does not fit this trend, but the higher rank 'positive' models do. As theoretically guaranteed [3], we observe no $\int \beta_1$ or $\int \beta_2$ for rank 1 for either model. For ranks 2-5, we observe that $\int \beta_1$ increases with an increase in matrix size for the 'mixed' model, but remains near 0 for the 'positive' model. A similar difference between the 'mixed' and 'positive' models is seen for $\int \beta_2$. **(b)** (Left) Wasserstein distances reproduced from Figure 5e for comparison. (Middle) Kullback-Leibler divergence of the distributions of $\int \beta_2$, $\int \beta_1$, and $\int \bar{\beta}_0$ for the random matrix models (columns) from the distributions for data-derived assembly correlations (rows). (Right) Jensen-Shannon distances between the distributions. We observe similar trends using the KL divergence and JS distance as with the Wasserstein distance. The KL divergence and JS distance have the drawback that they require binning the distributions of integrated Betti values. For each $\int \bar{\beta}_0, \int \beta_1$, and $\int \beta_2$, we use 20 bins of equal widths for the ranges $[0.05, 0.35], [0, 1.5], [0, 0.6]$, respectively. Another drawback of the KL divergence is that it can be infinite (black) when the model distribution has an empty bin and the data distribution does not, and it is not symmetric.

**Supplementary Figure 7. Sorted singular values of assembly correlations.** We consider assemblies with at least 20 neurons and use the dot product between the smoothed calcium signals of neurons as our measure of correlation. **(a)** Sorted singular values of assembly correlations during spontaneous activation ('on') for each assembly (gray curves), as well as the average largest 20 sorted singular values across assemblies (black), one plot per larva. We omit the largest singular value and normalize by dividing by the second-largest singular value. **(b-d)** Same as in (a), but for assembly correlations across the entire hour long recording ('whole'), outside of spontaneous activation ('off'), and for entrywise shuffles of the correlation matrices for the entire recording ('shuffle').

**Supplementary Figure 8. Additional low rank models and their Betti curves. (a)** Average Betti curves $\beta_0$ (gray), $\beta_1$ (blue), and $\beta_2$ (red) for (N = 100) size 50 matrices from random low rank matrix models for ranks $d =$ 1-5, where matrix entries $A_{ij} = \langle p_i, p_j \rangle$ for $p_i, p_j$ are drawn uniformly at random from different subsets of $\mathbb{R}^d$ for each of the 'mixed', 'positive', and 'positive + negative' models: $[-1, 1]^d$, $[0, 1]^d$ and $[-1, 0]^d \cup [0, 1]^d$, respectively. Notice that the 'positive' and 'positive + negative' models have similar Betti curves (top row). Another variation of low rank matrix models takes the negation of the matrices, $A_{ij} = -\langle p_i, p_j \rangle$. The 'negated' low rank matrix models produce very distinct Betti curves in each of the 'positive', 'mixed', and 'positive + negative' scenarios (bottom row). In particular, computing Betti curves for a negated correlation matrix can distinguish the 'positive' and 'positive + negative' models, even though the usual Betti curves do not. **(b)** Average Betti curves $\beta_0$ (gray), $\beta_1$ (blue), and $\beta_2$ (red) for *negated* assembly correlation matrices across the entire recording (N = 450 assemblies, 'whole'), during activations (N = 435 assemblies, 'on'), outside of activations (N

= 435 assemblies, 'off'), and for randomized assemblies across the entire recording (N = 450 assemblies, 'randomized assemblies'). Although many of the original Betti curves for these assembly correlation matrices were consistent with all three models, the Betti curves of the negated correlations are only consistent with the 'positive' model. In general, considering Betti curves for both the correlation matrix, $C_{ij}$, and its negated version, $-C_{ij}$, can reveal additional topological structure that enables the disambiguation of distinct underlying models.

**Supplementary Figure 9. Integrated Betti time series are not affected by the choice of sliding window size, and are similar across larvae. (a)** To check for the effects of sliding window size on the integrated Betti time series $\int \bar{\beta}_0(t)$ and $\int \beta_1(t)$ around activation onset, we computed the average and standard error for onset-aligned $\int \bar{\beta}_0(t)$ and $\int \beta_1(t)$ (N = 5046 activations across 12 larvae) for sliding windows of sizes $\approx$ 0.06 seconds (1 time bin), 0.25 seconds (4 time bins), 0.5 seconds (8 time bins, boxed in red), 1.0 seconds (15 time bins), and 2.0 seconds (30 time bins); all show similar trends. $\int \bar{\beta}_0(t)$ increases at onset while $\int \beta_1(t)$ crashes to zero at onset. All sliding window computations use a step of size $\approx$ 0.06 seconds (1 time bin). **(b)** Mean and standard error for the onset-aligned $\int \bar{\beta}_0(t)$ and $\int \beta_1(t)$ using a sliding window size 0.5 seconds (8 time bins) for each larva. (Top) Curves reproduced from Figure 6b for comparison. The overall average across al 12 larvae shows that $\int \bar{\beta}_0(t)$ increases at onset while $\int \beta_1(t)$ crashes to zero at onset. (Bottom) Similar trends hold for the individual larvae.

**Supplementary Figure 10. Determining the number of k-means clusters. (a)** To help determine the choice of $k$ for our k-means clustering of assembly activations from their $\int \bar{\beta}_0(t)$ time series, we performed a sillhouette analysis on the k-means clusters of $\int \bar{\beta}_0(t)$ (N = 5046 activations across 12 larvae) for $k = 2, \ldots, 12$ clusters. We observed that the average sillhouette score achieves a plateau centered at $k = 6$ before decreasing further. This motivated using $k = 6$ for our subsequent analyses. **(b)** We show the $\int \bar{\beta}_0(t)$ in each k-means cluster (color) and the average $\int \bar{\beta}_0(t)$ for each cluster (black) for $k = 3, 4, 5, 6, 7$ and 12. As we increase $k$, we observe smaller clusters emerging with distinct average $\int \bar{\beta}_0$ profiles. For $k = 12$, on the other hand, these distinctions are less pronounced, suggesting an over-refined partition of the activations. For each k-means clustering, we used the segment of $\int \bar{\beta}_0(t)$ that is $[-0.3, 0.67]$ seconds around onset.

**Supplementary Figure 11. Effects of choice of time window on k-means clusters.** Here we checked the effect of the choice of temporal window of $\int \bar{\beta}_0(t)$ used to perform the k-means clustering of the assembly activations (N = 5046 activations across 12 larvae), for fixed $k = 6$. **(a)** We show the 6 clusters obtained when using the entire $[-2s, 2s]$ time window around activation onset. Note that the clusters are distinguished by timing and magnitude of the rise in $\int \bar{\beta}_0$ relative to activation onset. **(b-f)** Same as in (a), but for time windows $[-1, 1], [-2, 0], [-1, 0], [0, 2]$, and $[0, 1]$ seconds around onset. The windows centered around the onset (a-b) yield similar clusters. For windows entirely pre-onset (c-d) or post-onset (e-f), the clusters emphasize somewhat different features, capturing distinctions between pre-onset and post-onset profiles, respectively. For the analyses in Figures 7 and 8, we chose a time window of $[-1/3, 2/3]$ seconds around activation onset.

**Supplementary Figure 12. Activations and cholinergic activity.** (Top) We reproduce Figure 7a showing the six k-means clusters of activation $\int \beta_0(t)$ profiles across all activations (N=5046) of assemblies across 12 larvae. (Bottom) Each row shows the timing of all assembly spontaneous activations per larva. Each dot (colored by cluster membership) shows an activation of an assembly. Below we show the average total neural activity (gray) and average total cholinergic activity (blue) across the optic tectum per larva, using the smoothed calcium signal to represent neural activity. Note: the immunostaining in the larva CTR06 (top row) only captured cholinergic neurons in the first of three recorded layers.

### References


[1] Martin Rosvall and Carl T Bergstrom. Maps of random walks on complex networks reveal community structure, 2008.

[2] Gábor Csárdi and Tamás Nepusz. The igraph software package for complex network research. *InterJournal*, page 1695, 2006.

[3] Carina Curto, Joshua Paik, and Igor Rivin. Betti curves of rank one symmetric matrices. *Geometric Science of Information*, pages 645–655, 2021.

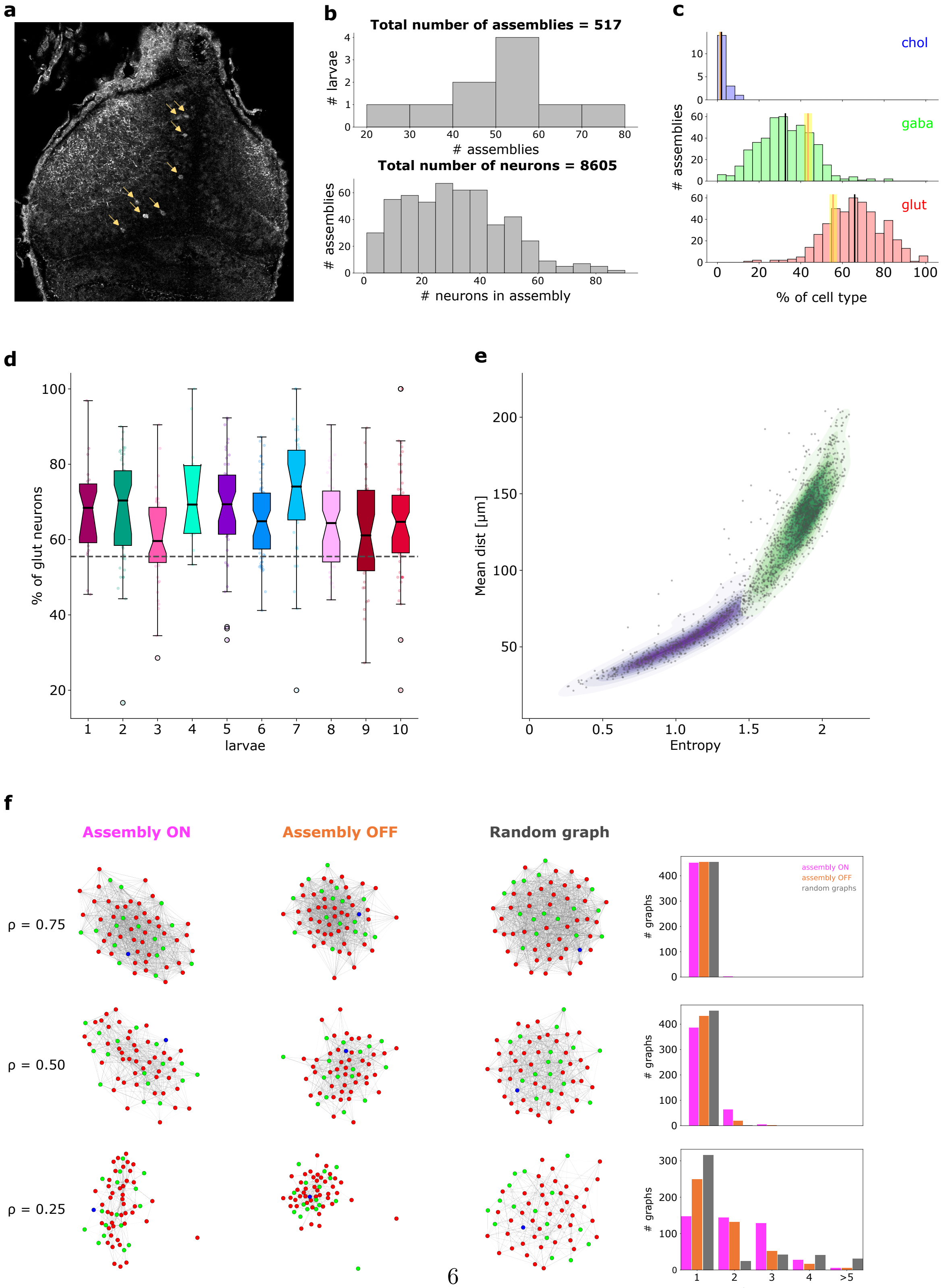




Figure 1: **Supplementary Figure 1** – see legend

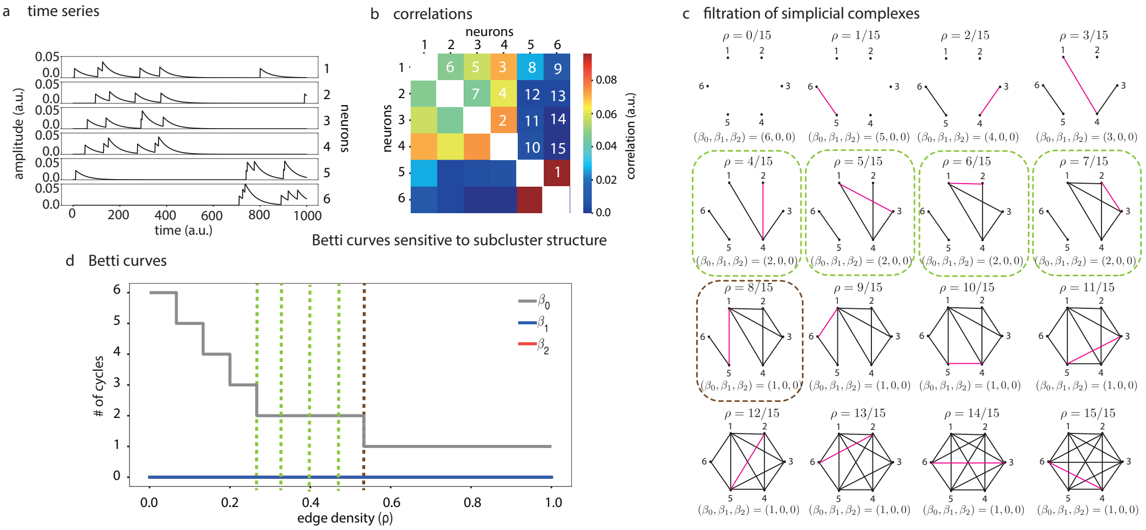


Figure 2: **Supplementary Figure 2** – see legend

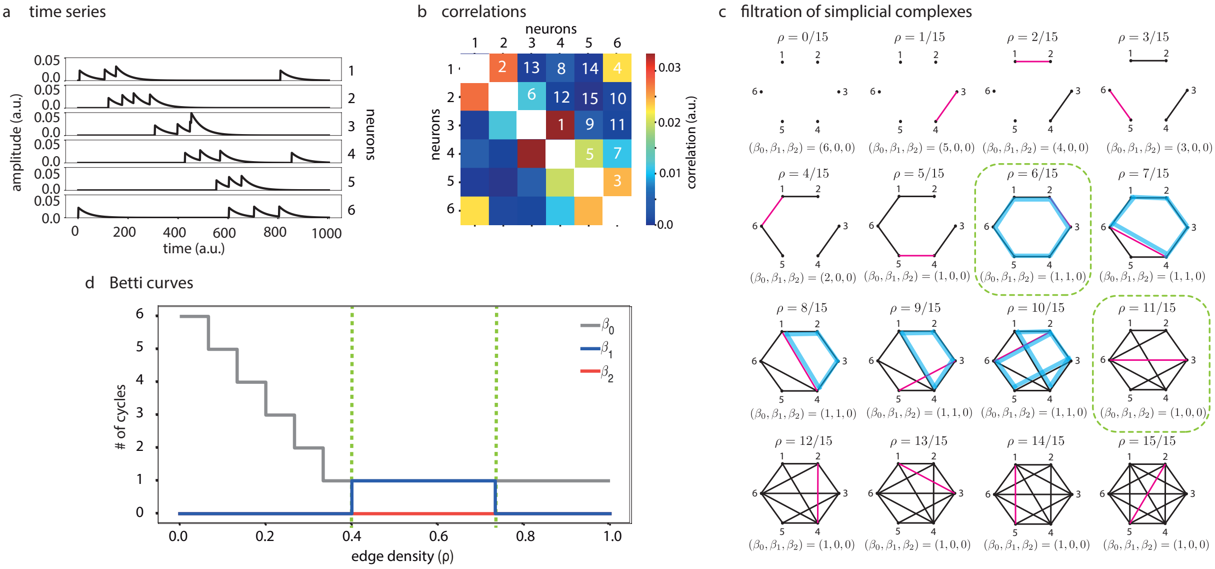


Figure 3: **Supplementary Figure 3** – see legend

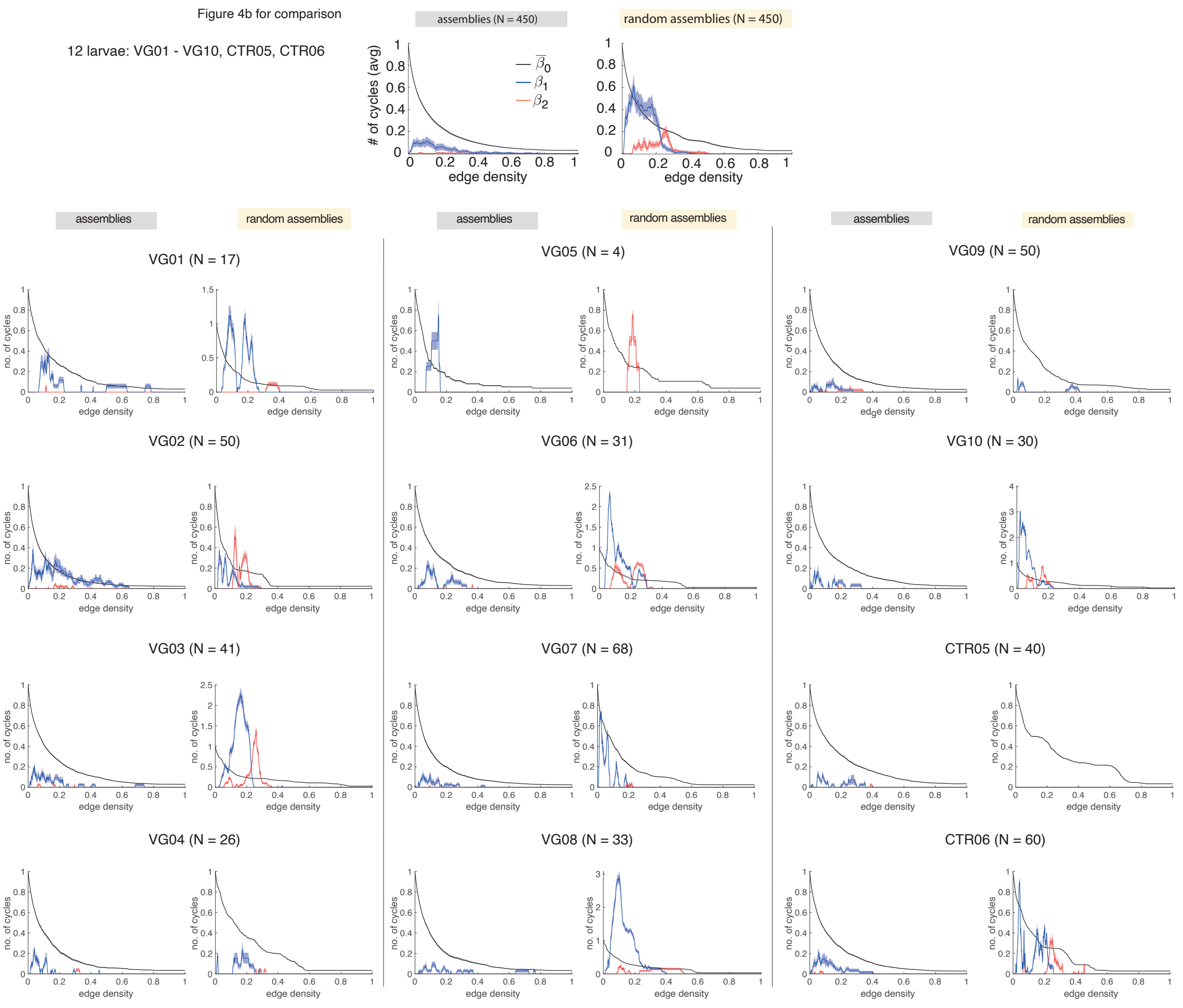


Figure 4: **Supplementary Figure 4** – see legend

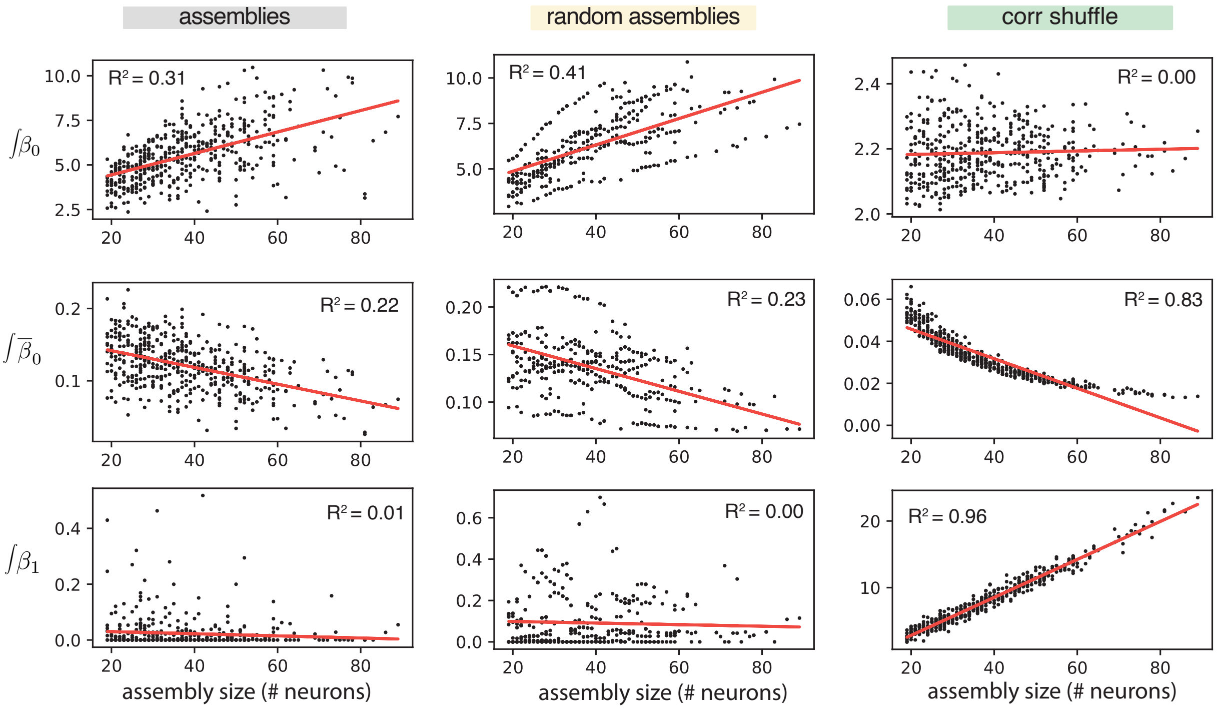


Figure 5: **Supplementary Figure 5** – see legend

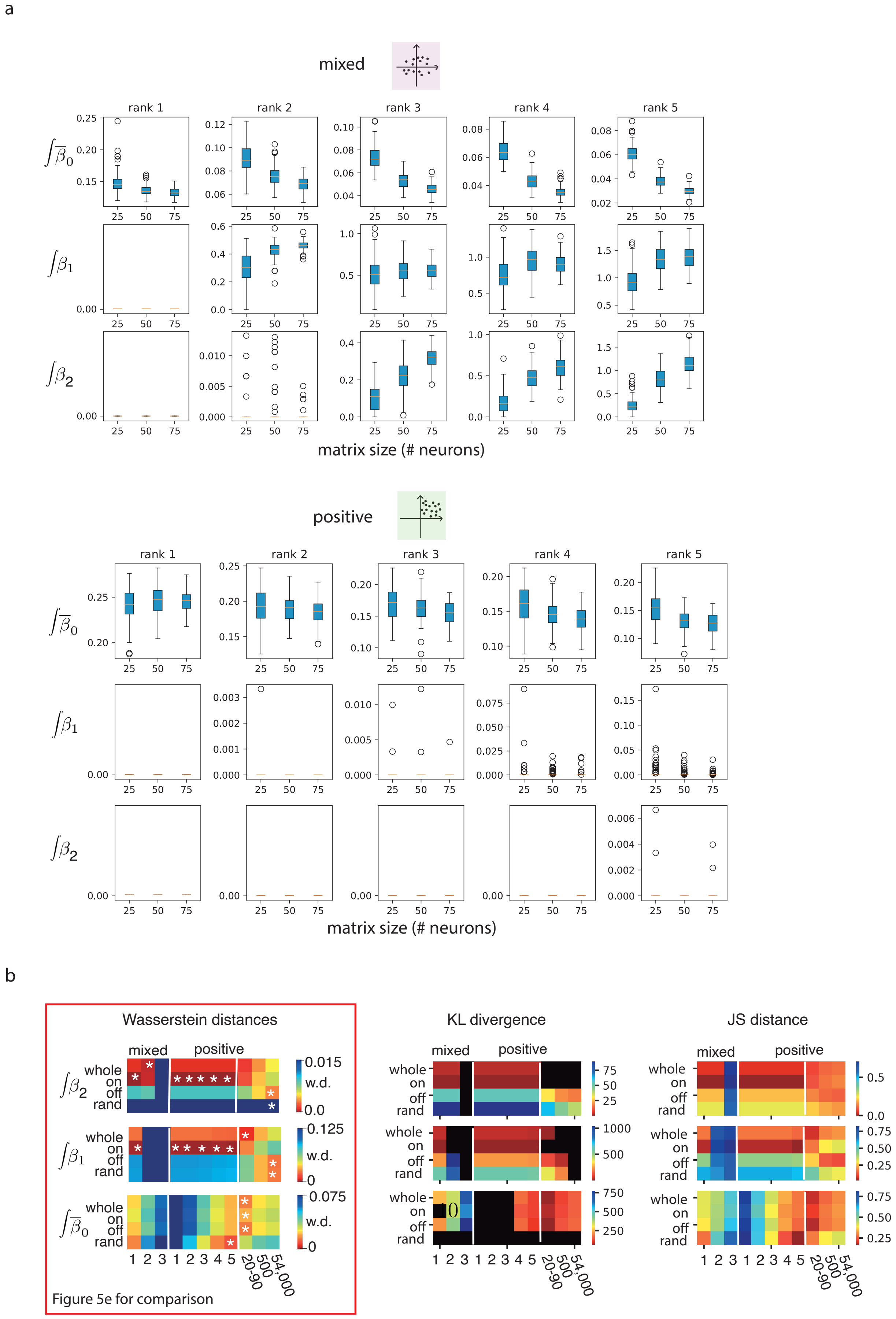


Figure 6: **Supplementary Figure 6** – see legend

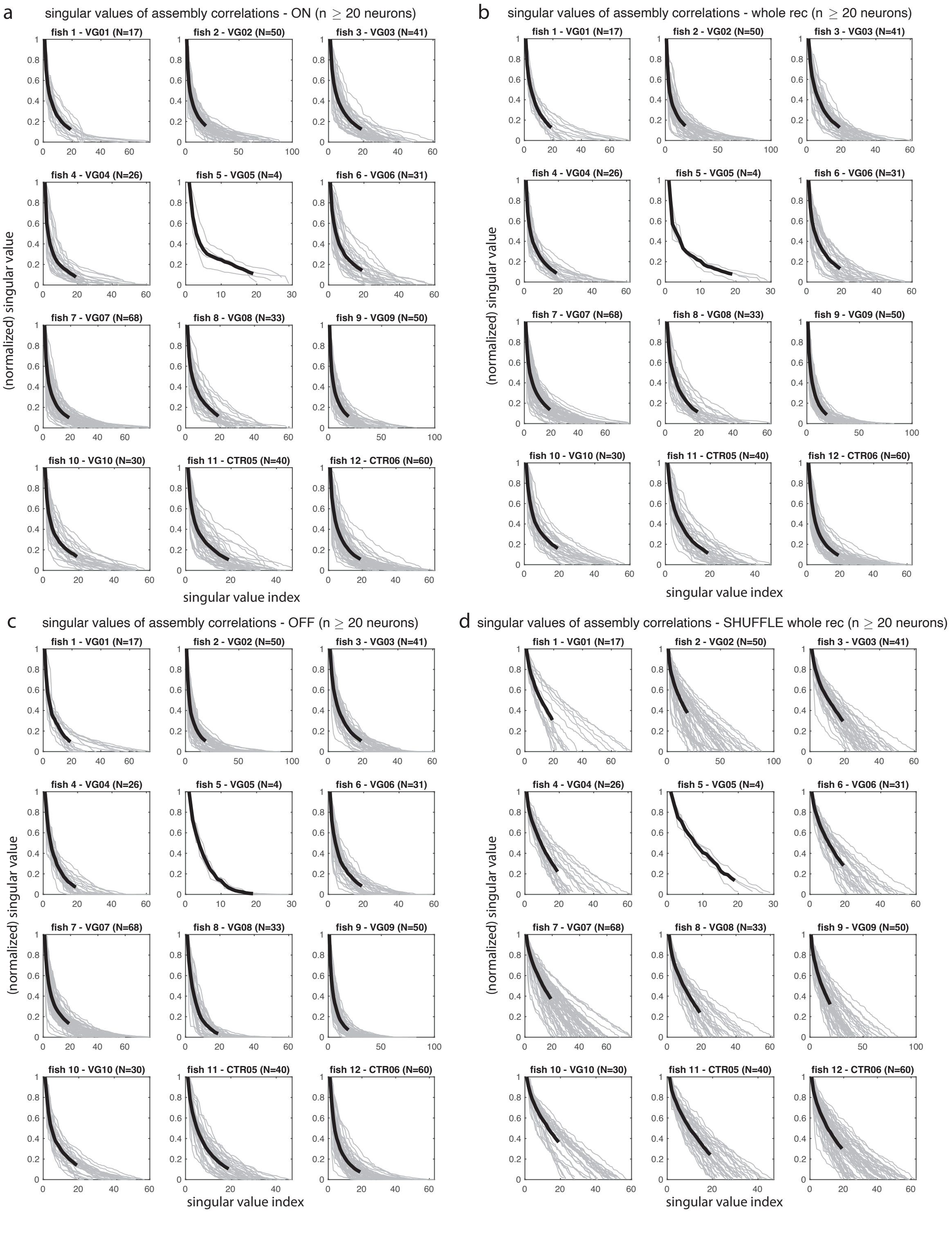


Figure 7: **Supplementary Figure 7** – see legend

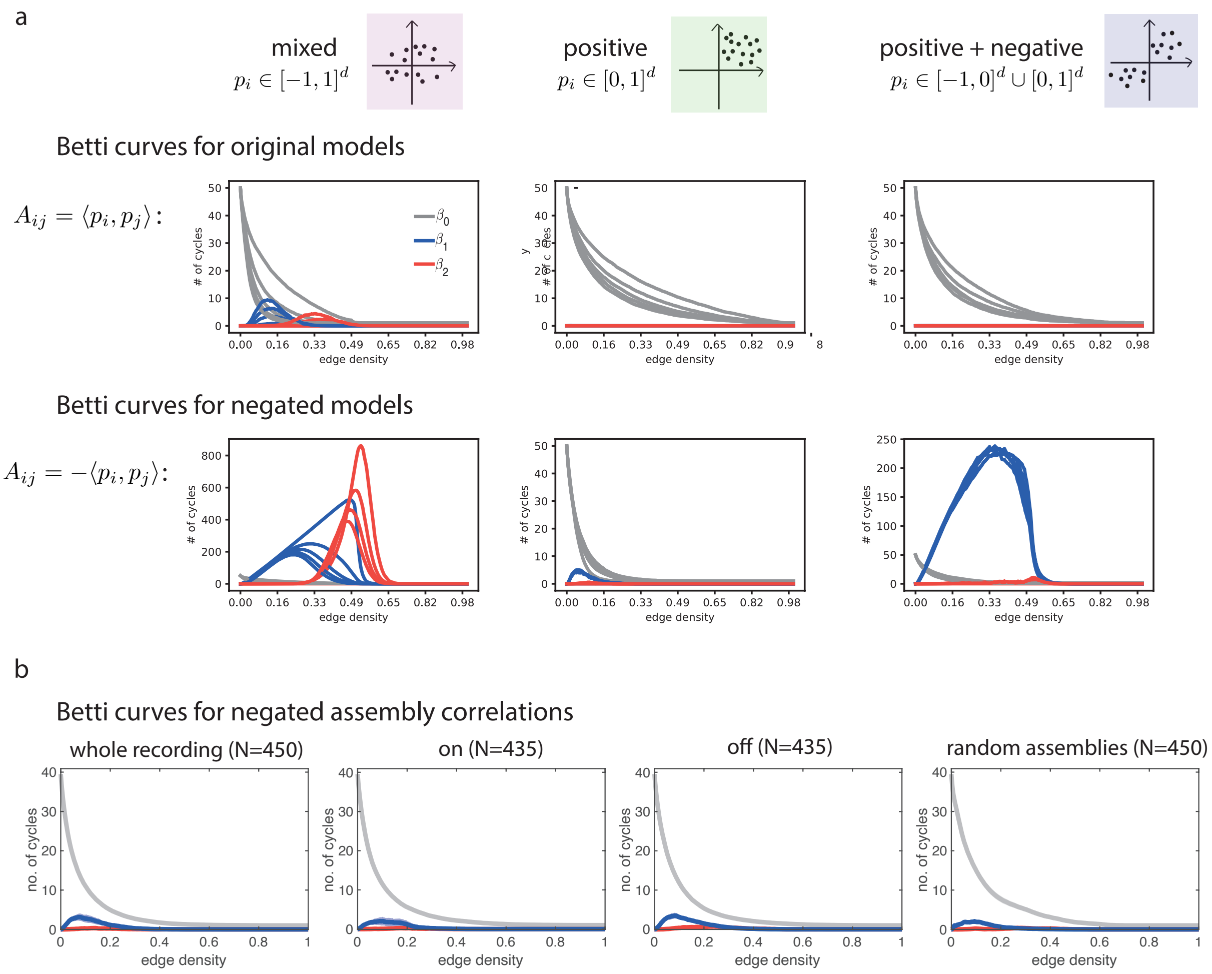


Figure 8: **Supplementary Figure 8** – see legend

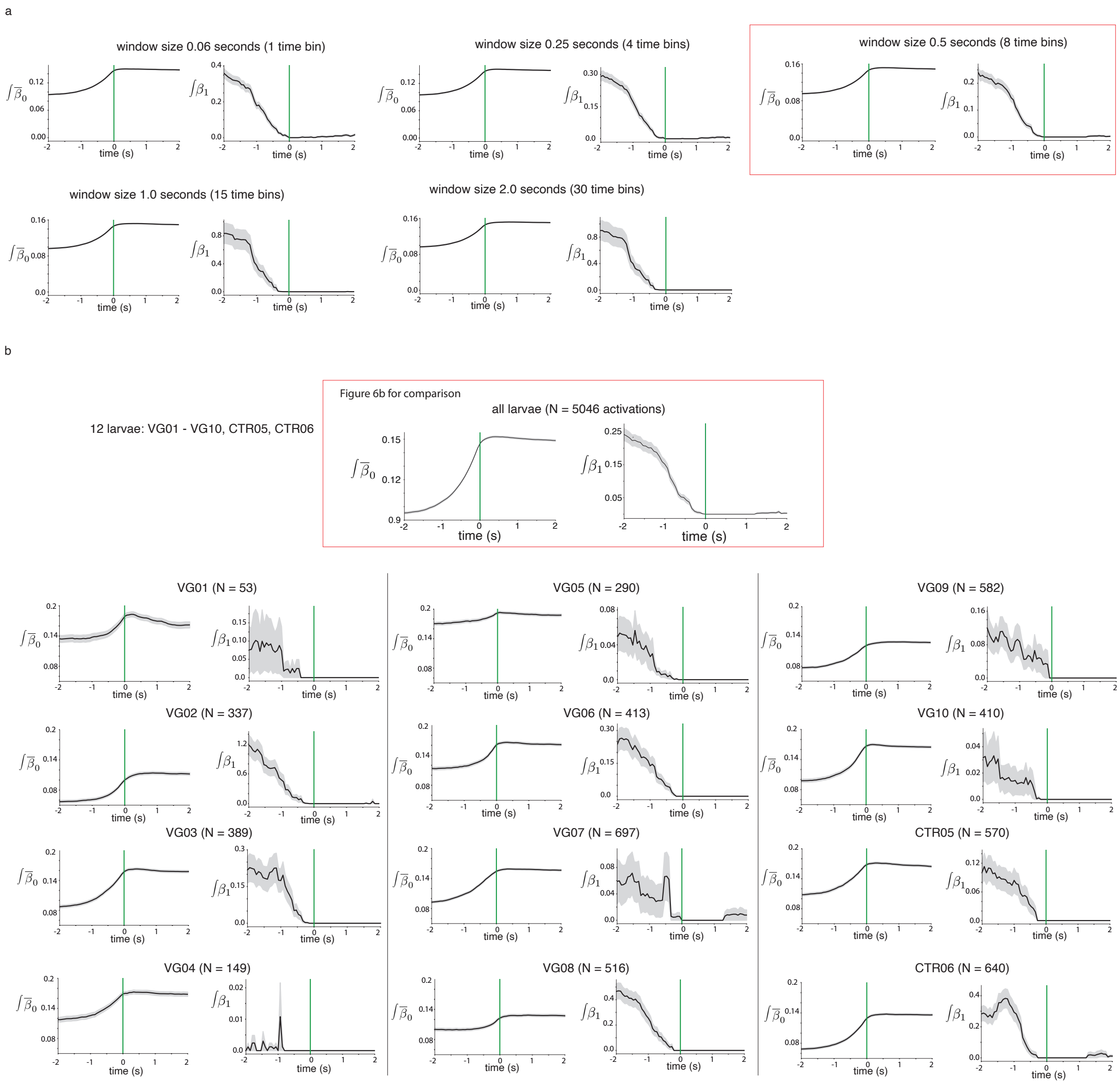

Figure 9: **Supplementary Figure 9** – see legend

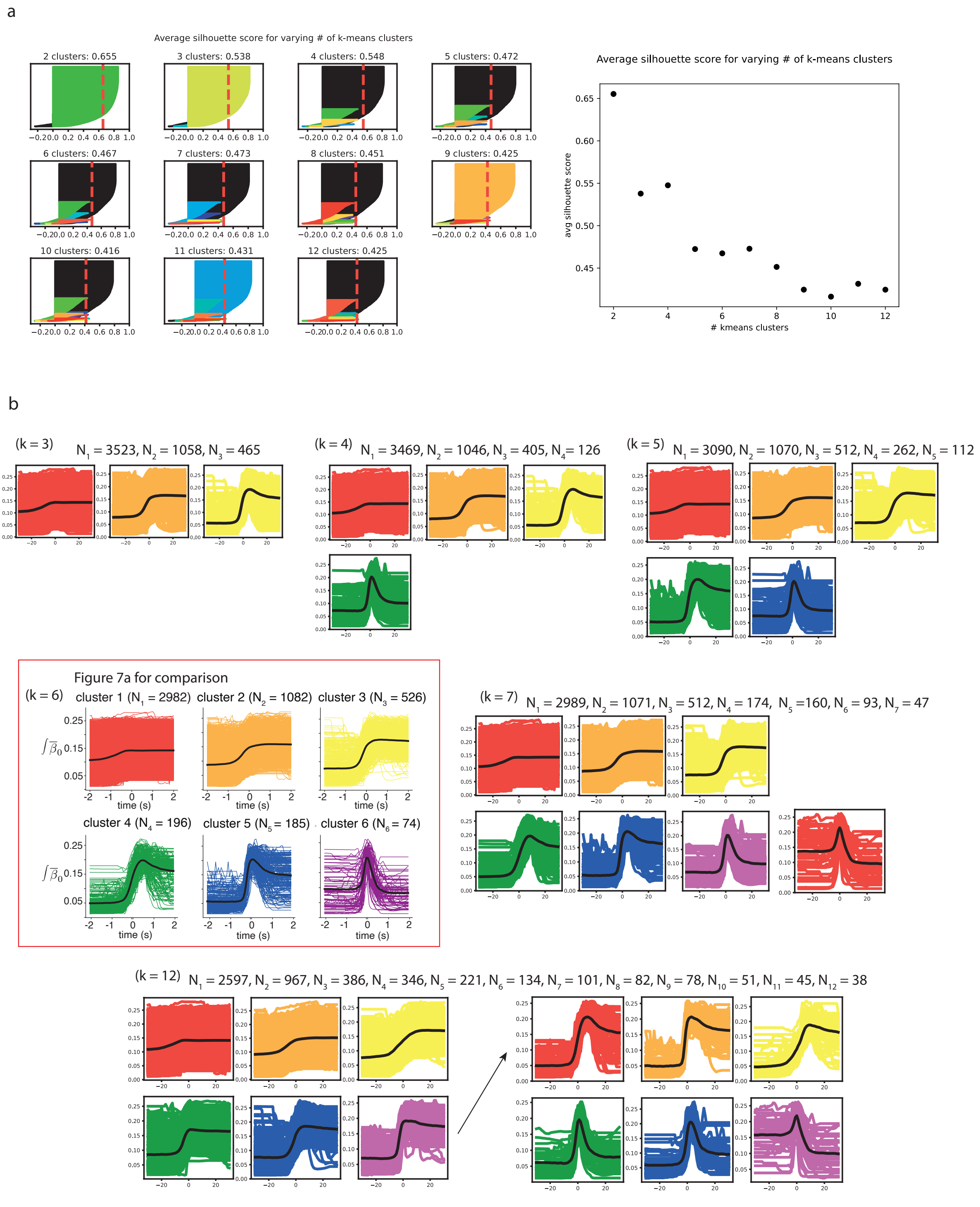


Figure 10: **Supplementary Figure 10** – see legend

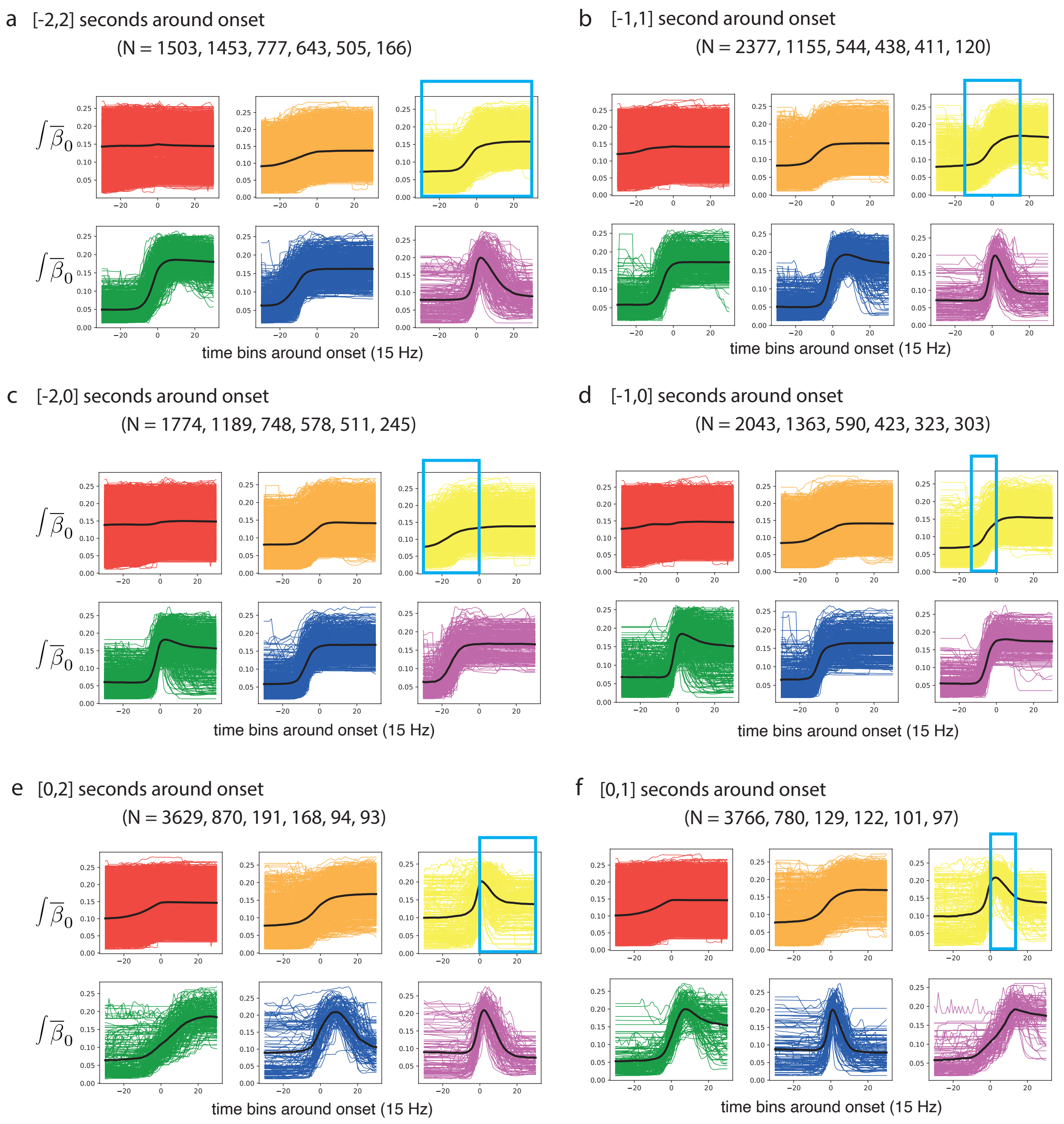


Figure 11: **Supplementary Figure 11** – see legend

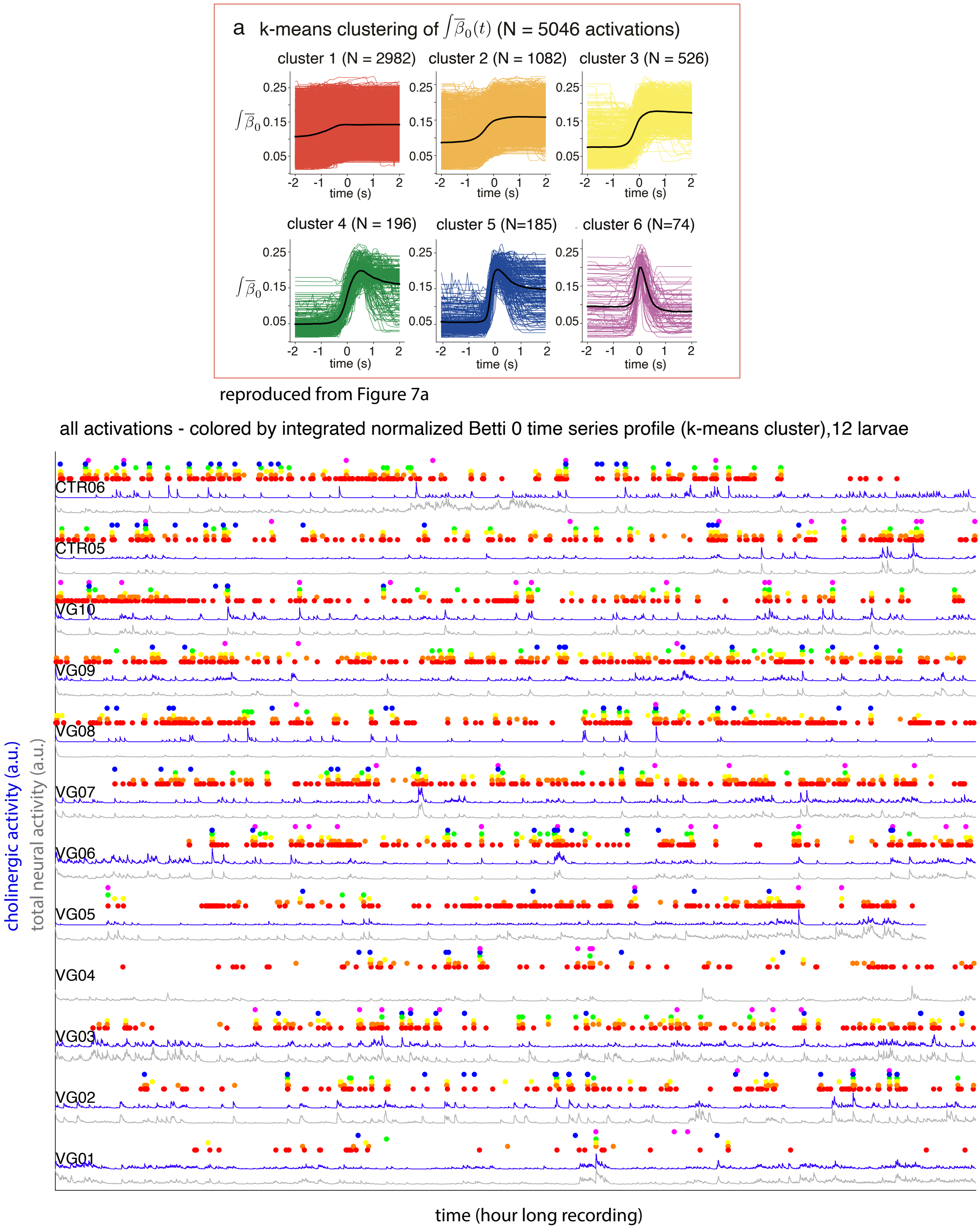


Figure 12: **Supplementary Figure 12** – see legend
